# XLF acts as a flexible connector during non-homologous end joining

**DOI:** 10.1101/2020.07.29.227033

**Authors:** Sean M. Carney, Andrew T. Moreno, Sadie C. Piatt, Metztli Cisneros-Aguirre, Felicia Wednesday Lopezcolorado, Jeremy M. Stark, Joseph J. Loparo

## Abstract

Non-homologous end joining (NHEJ) is the predominant pathway that repairs DNA double strand breaks in vertebrates. During NHEJ DNA ends are held together by a multi-protein synaptic complex until they are ligated. Here we investigate the role of the intrinsically disordered C-terminal tail of XLF, a critical factor in end synapsis. We demonstrate that the XLF tail along with the Ku binding motif (KBM) at the extreme C-terminus are required for end joining. While the underlying sequence of the tail can be varied, a minimal tail length is required for NHEJ. Single-molecule FRET experiments that observe end synapsis in real-time show that this defect is due to a failure to closely align DNA ends. Our data supports a model in which a single C-terminal tail tethers XLF to Ku while allowing XLF to form interactions with XRCC4 that enable synaptic complex formation.

## Introduction

DNA double strand breaks (DSBs) are a particularly toxic form of DNA damage. Within vertebrates the majority of DSBs are repaired by non-homologous end joining (NHEJ).^1^ In contrast to homologous recombination (HR), the other major DSB repair pathway, NHEJ does not use a DNA template to guide repair. Instead a synaptic complex comprised of a number of core and accessory NHEJ factors holds DNA ends together until they are ultimately ligated. Recognition of the DSB is carried out by the ring-shaped Ku70/Ku80 heterodimer (Ku) which rapidly binds DNA ends^2^ and subsequently recruits downstream NHEJ factors including the DNA-dependent protein kinase catalytic subunit, DNA-PKcs,^3,4^ whose kinase activity is essential for DNA repair.^5^ As DSBs arise from a wide-range of sources, DNA ends are often initially incompatible with ligation. A host of end processing enzymes, including NHEJ associated polymerases and nucleases, act on these ends to allow for ligation by DNA ligase IV (Lig4).^6,7^ Ligation requires at least two additional factors: XRCC4, a scaffolding factor to which Lig4 is constitutively bound, and the structurally related XRCC4-like factor (XLF).^6,8,9^ Together these factors must assemble into a synaptic complex that recognizes, synapses, aligns, processes, and ligates DNA ends.

The NHEJ synaptic complex holds DNA ends together through a complicated network of intermolecular interactions. Emerging single-molecule approaches in cell-free extracts and reconstitutions have provided new mechanistic details of how these interactions evolve during repair reactions. Using single-molecule Förster resonance energy transfer (smFRET) experiments to monitor the distance between DNA ends in *Xenopus laevis* egg extract, we have shown that there are at least two distinct synaptic states that precede ligation.^10^ First, Ku and DNA-PKcs (but not kinase activity) are required to weakly tether DNA ends at a distance where they are protected from processing, a state we named the long-range (LR) complex.^7,10^ Next, DNA-PKcs kinase activity, XRCC4, Lig4, and XLF are required to transition from the initial LR complex to a stable short-range (SR) synaptic complex in which the DNA ends are closely aligned for processing and ligation.^7,10^ Importantly, the catalytic activity of Lig4 is not required to form the SR complex, demonstrating that the ligase plays a structural role in end synapsis.^10,11^ Subsequent biochemical reconstitutions of human NHEJ proteins also found evidence for these two synaptic states,^12^ suggesting that the architecture of the NHEJ synaptic complex is conserved from *Xenopus* to humans.

The role of XLF in end synapsis remains especially vague as it has no known catalytic activity, and conflicting models of its function have obscured its role in NHEJ. XLF exists as a homodimer along with two other NHEJ proteins, the structurally similar XRCC4 and the NHEJ accessory factor, PAXX.^13,14,15^ Each of these paralogues consists of a N-terminal globular head domain, an extended coiled-coil stalk that mediates dimerization,^15^ and a flexible C-terminal tail (FIG 1A).^16^ XLF and XRCC4 interact through their head domains^17^,^18^ and this interaction is essential for NHEJ in cells.^19,20^ In minimal reconstitutions of XRCC4 and XLF, this interaction can lead to the formation of extensive XLF-XRCC4 filaments^17,21^ which have been proposed to be important for repair *in vivo*.^22,23^ However, single-molecule imaging of fluorescently labeled XLF in egg extract by our laboratory revealed that a single XLF dimer is sufficient for SR complex formation, although it has to be able to interact with XRRC4 through both of its head domains for optimal end joining.^24^ These results suggest that a single XLF dimer makes several contacts within the SR complex to mediate synapsis.

**Figure 1.**
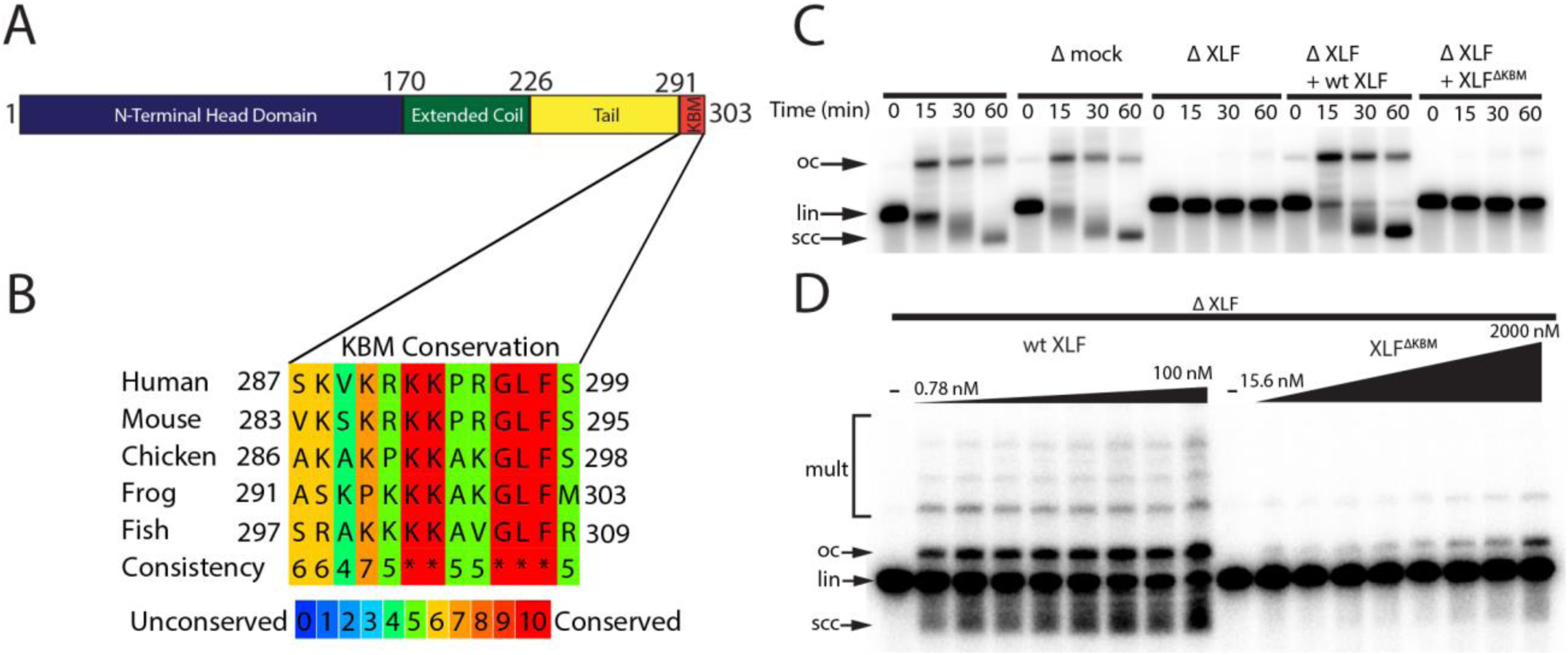
The Ku Binding Motif (KBM) of XLF is critical for end joining in *Xenopus* egg extract. **A**, A schematic of XLF’s domain organization. The residue number at the boundary of each region is shown. **B**, A protein sequence alignment of the KBM from human (*Homo sapiens* UniProt ID Q9H9Q4), mouse (*Mus musculus* UniProt ID Q3KNJ2), chicken (*Gallus gallus* UniProt ID F1NVP8), frog (*Xenopus laevis* see note below), and fish (*Danio rerio* UniProt ID Q6NV18). The *Xenopus laevis* XLF sequence and translation start site was determined previously by immunoprecipitation from extract and subsequent trypsin digestion and analysis by mass spectrometry.^10^ The alignment was performed using PRALINE multiple sequence alignment and its default settings.^43^ A colored scale shows the degree of conservation. **C**, Ensemble time course end joining assay in either mock-depleted (immunodepletion with non-specific rabbit IgG) or XLF-depleted *Xenopus* egg extract. Recombinant wild type (wt) XLF and XLF^ΔKBM^ were added back at 75 nM final concentration. DNA species: scc, supercoiled closed circular; lin, linear; oc, open circle. **D**, Ensemble end point titration end joining assay in *Xenopus* egg extract. XLF-depleted extract was supplemented with recombinant protein, either wt XLF or XLF^ΔKBM^, at varying concentrations. The reactions were stopped after 20 minutes. DNA species: scc, supercoiled closed circular; lin, linear; oc, open circle; mult, multimer.

XLF contains an extended intrinsically disordered C-terminal tail with a highly conserved Ku binding motif (KBM) at the terminus (FIG 1 A-B).^25,26^ Deletion of the KBM ablates cellular recruitment of XLF to sites of DNA damage^25^ and mutations within the KBM result in varying defects in cellular end joining assays and cell survival in the presence of DSB inducing agents.^26,27^ The contributions of the remainder of the C-terminal tail, henceforth referred to as the tail, are unknown, although it has been shown to be a target for phosphorylation by DNA-PKcs.^28^ Across vertebrate species, there is poor sequence conservation within this region, and yet the tail is invariably retained with a similar length (Supp. FIG 1A). Deleting the entire C-terminal region of XLF (including the KBM) still allows for interaction with XRCC4.^16^ However, minimal reconstitutions show that the C-terminal region of XLF is necessary for an interaction with DNA *in vitro*^16,17^ and variably stimulates (1.5-40 fold) end joining.^12,16^ It remains unclear if these defects arise solely from the loss of the XLF KBM or if the tail also contributes to NHEJ.

Here, we investigate the role of the XLF C-terminal tail in NHEJ synaptic complex assembly. We show that both the KBM and the tail are essential for robust end joining in both *Xenopus* egg extract and in cells. Ablation of potential phosphorylation sites and shuffling the sequence of the tail region did not alter repair efficiency, demonstrating that, independent of sequence, the length of this region is important for NHEJ. Using asymmetric mutants of XLF, we show that a single KBM within the XLF dimer is sufficient for end joining. These observations rule out models where both XLF tails are required to span the break via interactions with opposing Ku molecules. Instead, the length of the tail is required for XLF-mediated stabilization of XRCC4-Lig4 at DNA ends. We propose that the XLF tail acts to tether XLF to DSBs through its interaction with Ku while simultaneously allowing XLF to interact with XRCC4 and drive formation of the SR complex.

## Results

### The Ku Binding Motif (KBM) of XLF is essential for end joining in *Xenopus* egg extract

To test whether the KBM of XLF is necessary for end joining, we performed a time course end joining assay in *Xenopus* egg extract. In this assay, we monitored the conversion of radio-labeled linear DNA to joined products (open circular DNA, supercoiled closed DNA, and in some cases multimers) over time (FIG 1C). Immunodepletion of XLF from extract (Supp. FIG 1B) abolished end joining (FIG 1C), consistent with prior results in extract and in cells.^10,19^ A mock immunodepletion did not alter end joining, and joining in XLF-depleted extract could be rescued by adding back physiological concentrations (∼75 nM) of recombinant wild type (wt) XLF (FIG 1C).^29^ Notably, an XLF mutant lacking the KBM, XLF^ΔKBM^, did not rescue end joining in XLF-depleted extract, demonstrating that the KBM is necessary for end joining. To further characterize the severity of this defect, we performed titrations of wt XLF and XLF^ΔKBM^ in XLF-depleted extract (FIG 1D). While wt XLF was able to robustly rescue joining at sub-nanomolar concentrations, the XLF^ΔKBM^ supported little end joining even at micromolar concentrations. Collectively these results are consistent with the KBM playing a critical role in recruiting XLF to DSBs through its interaction with Ku.^25,26^

### The interior region of the C-terminal tail of XLF is essential for end joining

Given the importance of the KBM, we next tested whether the tail of XLF is also required for end joining. We constructed several mutants in which the tail was truncated while retaining the KBM at the very C-terminus (FIG 2A). The three truncation constructs XLF^1-285+KBM^, XLF^1-265+KBM^, and XLF^1-245+KBM^ removed 6, 26, and 46 residues respectively from the tail (FIG 2A). To determine if the unstructured tail of XLF is necessary for NHEJ, we examined the ability of each C-terminal truncation mutant to rescue end joining in XLF-depleted extract (FIG 2B). At 75 nM the more severe truncations, XLF^1-265+KBM^ and XLF^1-245+KBM^, showed no significant joining activity, while the most conservative truncation, XLF^1-285+KBM^, was able to rescue end joining. Even at a concentration of 500 nM, joining by XLF^1-245+KBM^ was barely detectable while joining by XLF^1-265+KBM^ was clearly much slower than the wild type. In contrast, joining by XLF^1-285+KBM^ was similar to the wild type (Supp. FIG 2A). To ensure that these defects in end joining did not arise due to protein misfolding, we measured the stability of each mutant using differential scanning fluorimetry (Supp. FIG 2B). Consistent with all three mutants being stably folded, we found that the melting temperatures (T_m_) of the mutants were similar to the T_m_ of wt XLF. Thus, the tail of XLF is required for efficient end joining in egg extract.

**Figure 2.**
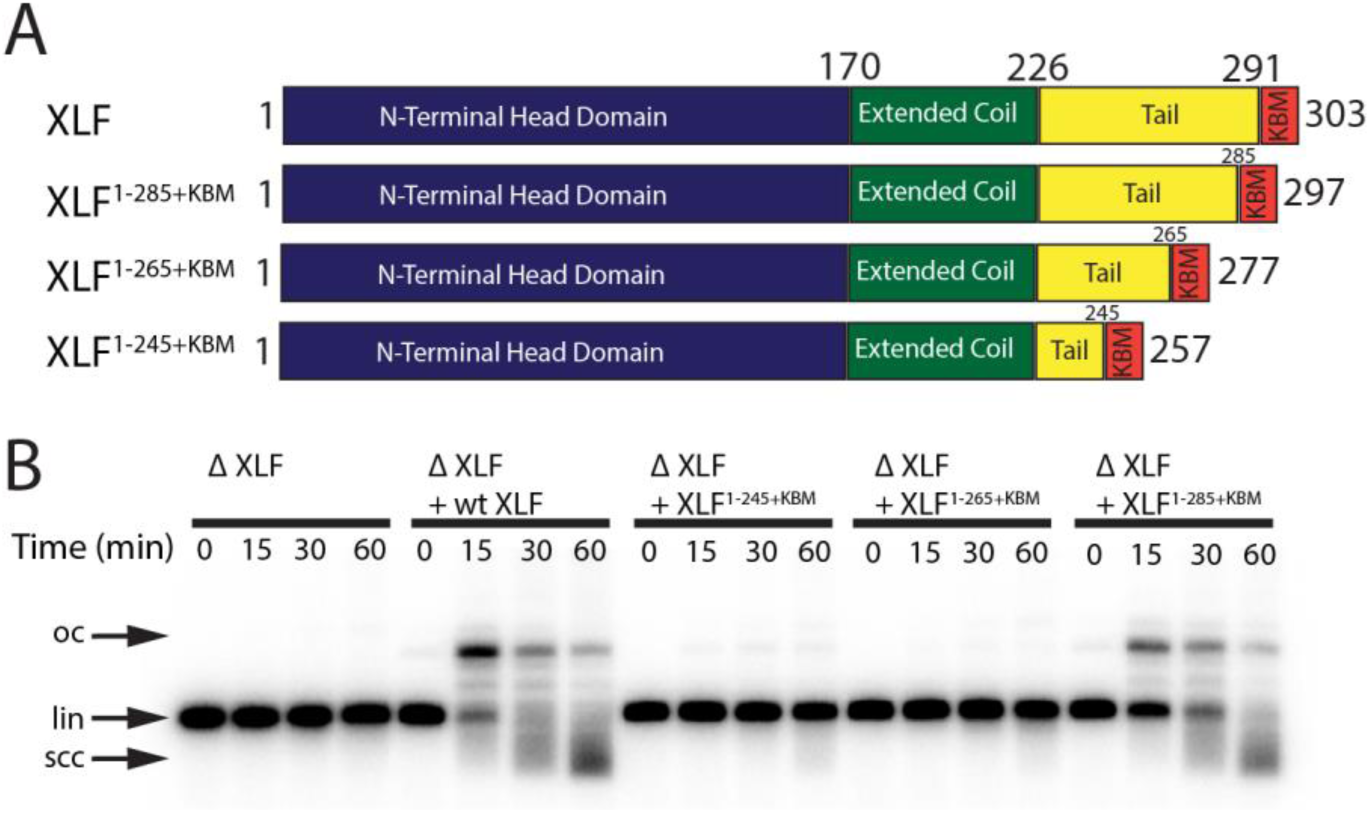
The tail region of XLF is essential for DNA end joining. **A**, Schematics of the domain organization of wt XLF and truncation mutants (XLF^1-285+KBM^, XLF^1-265+KBM^, and XLF^1-245+KBM^). The residue number at the boundary of each region of the protein is indicated. **B**, Ensemble time course end joining assay in XLF-depleted *Xenopus* egg extract. Recombinant wt XLF, XLF^1-285+KBM^, XLF^1-265+KBM^, and XLF^1-245+KBM^ were added back at 75 nM final concentration to their respective reaction samples. DNA species: scc, supercoiled closed circular; lin, linear; oc, open circle.

### The interior region of the XLF C-terminal tail is required to form short-range synaptic complex

To further interrogate how truncating the tail of XLF leads to defects in end joining, we used a smFRET assay that reports on the formation of the SR complex. This assay utilizes a 2 kb linear DNA labeled with Cy3 and Cy5 dyes 7 nt from each blunt end of the substrate and contains an internal biotin that is used to immobilize it on the surface of a flow cell (FIG 3A).^10,30^ FRET between Cy3 and Cy5 only occurs within the SR complex with the average FRET efficiency being indistinguishable from the ligated product.^10^ For these experiments, extract depleted of XLF was supplemented with either buffer or recombinant protein and flowed into the flow cell. Each replicate consisted of a movie of three fields of view (FOVs) imaged for 15 minutes each. Example trajectories from the ΔXLF + wt XLF condition are shown in FIG 3B-C. In these trajectories two distinct FRET states are observed: a low- or no-FRET state corresponding to unpaired DNA ends or the LR complex and a high-FRET state that corresponds to the SR complex and ultimately the ligated DNA product. FIG 3B shows an example trajectory where the SR complex forms and subsequently falls apart. FIG 3C shows an example where SR complex formation leads to the high-FRET state that persists until the end of the observation window. In FIG 3D-F the FRET signal is plotted as a normalized histogram for each 15-minute interval of the experiment. In the case of the ΔXLF + wt XLF condition, we observed a time-dependent increase in the high-FRET population (FRET efficiency ∼ 0.5) due to accumulation of the SR complex along with ligated products (FIG 3E). In contrast, the high-FRET population was not observed in the ΔXLF + buffer (FIG 3D) or ΔXLF + XLF^1-245+KBM^ (FIG 3F) conditions. The absence of a high-FRET population in the XLF^1-245+KBM^ condition could be due to (1) a substantial decrease in the stability of the SR complex or (2) an inability of the SR complex to form. To distinguish between these possibilities, we measured the rate of SR complex formation by recording the number of individual FRET events detected from all single molecules tracked (Supp. Table 1). The SR complex formation rate for wt XLF (1.7×10^−3^ ± 3.4×10^−4^ s^−1^) agrees well with a previously published value,^24^ while the rates for the buffer and XLF^1-245+KBM^ conditions were >25-fold lower, at 6.4×10^−5^ ± 1.0×10^−5^ s^−1^ and 4.6×10^−5^ ± 1.3×10^−5^ s^−1^, respectively (FIG 3G). As we observed a similarly low number of high FRET events (SR complex formation) for the buffer and XLF^1-245+KBM^ conditions (Supp. Table 1), these results show that XLF^1-245+KBM^ is deficient in forming the SR complex. Given the severe defect in SR complex formation rates for these conditions, we were unable to collect enough events to compare the stability of the SR complex. Collectively, these results indicate that the tail of XLF is required for formation of the SR complex and the close alignment of DNA ends.

**Figure 3.**
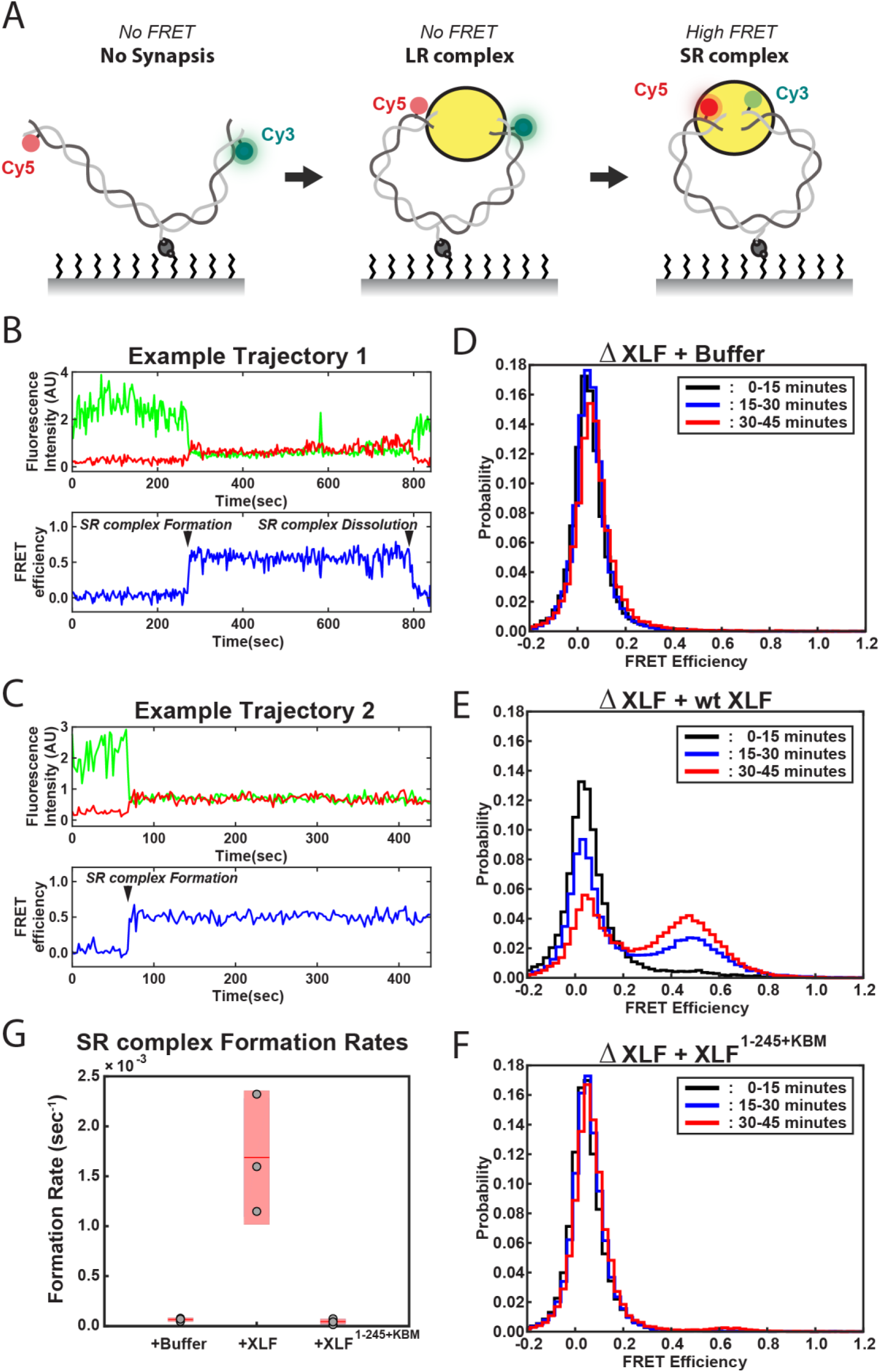
DNA end synapsis requires the tail of XLF. **A**, Schematic of the FRET-labeled DNA substrate immobilized in the flow cell via biotin-streptavidin interactions. In the absence of egg extract the DNA substrate exhibits no FRET. When the LR complex forms, the DNA ends are not positioned close enough together for energy transfer from Cy3 to Cy5 even though the ends are co-localized within the NHEJ synaptic complex.^10^ Upon formation of the SR complex, FRET between the fluorophores on opposing DNA ends can be observed. **B**,**C**, Example trajectories that contain SR complex formation events. Donor and acceptor fluorescence intensity are shown in green and red, respectively. The corresponding FRET efficiency from each trace is shown in blue in a separate trajectory below. **D, E, F**, Normalized FRET histograms for each experimental condition accumulated over a 15-minute time window. Data from each 15-minute field of view is represented by a separate curve. **G**, Plot of SR complex formation rates. For each condition, individual replicates are plotted as grey circles, and the mean is represented as dark red horizontal line. The 95% confidence interval is represented for each condition as a light red rectangle centered on the mean. This plot was generated using the notBoxPlot MATLAB function.^42^

### The sequence of the C-terminal tail of XLF is not critical for its role in end joining

The synapsis and joining defects seen for the XLF truncation mutants suggest two potential roles for the C-terminal tail of XLF: (1) the length of the XLF tail may be critical within the architecture of the NHEJ synaptic complex, allowing XLF to interact with binding partners Ku80 and XRCC4; alternatively, (2) residues within the C-terminal tail may be critical sites of phosphorylation or make contacts with other factors. Both DNA-PKcs and ATM are known to target sites within the XLF tail for phosphorylation.^28^ To test whether phosphorylation sites within the tail are required for end joining, we mutated all serines to glycines and all threonines to alanines between residues 226 and 292. This mutant, XLF^NoPhos+KBM^, was able to rescue end joining in XLF-depleted extract as efficiently as wt XLF (FIG 4A), consistent with XLF phosphorylation not being required for end synapsis and ligation. To disrupt any unidentified interaction motifs, we randomly shuffled the sequence between residues 226 and 292 of XLF (Supp. FIG 3A). Sequences were shuffled using the Sequence Manipulation Suite while the KBM was left unaltered since it is necessary for end joining (FIG 1C).^31^ We constructed two distinct shuffled mutants using this approach, XLF^ShuffA+KBM^ and XLF^ShuffB+KBM^ (Supp. FIG 3A), and each of these mutants rescued end joining in XLF-depleted extract with wild type-like efficiency (FIG 4A). Collectively these results rule out the loss of phosphorylation sites or disruption of a required motif within the tail as the cause of the end joining defect exhibited by the truncations mutants.

**Figure 4.**
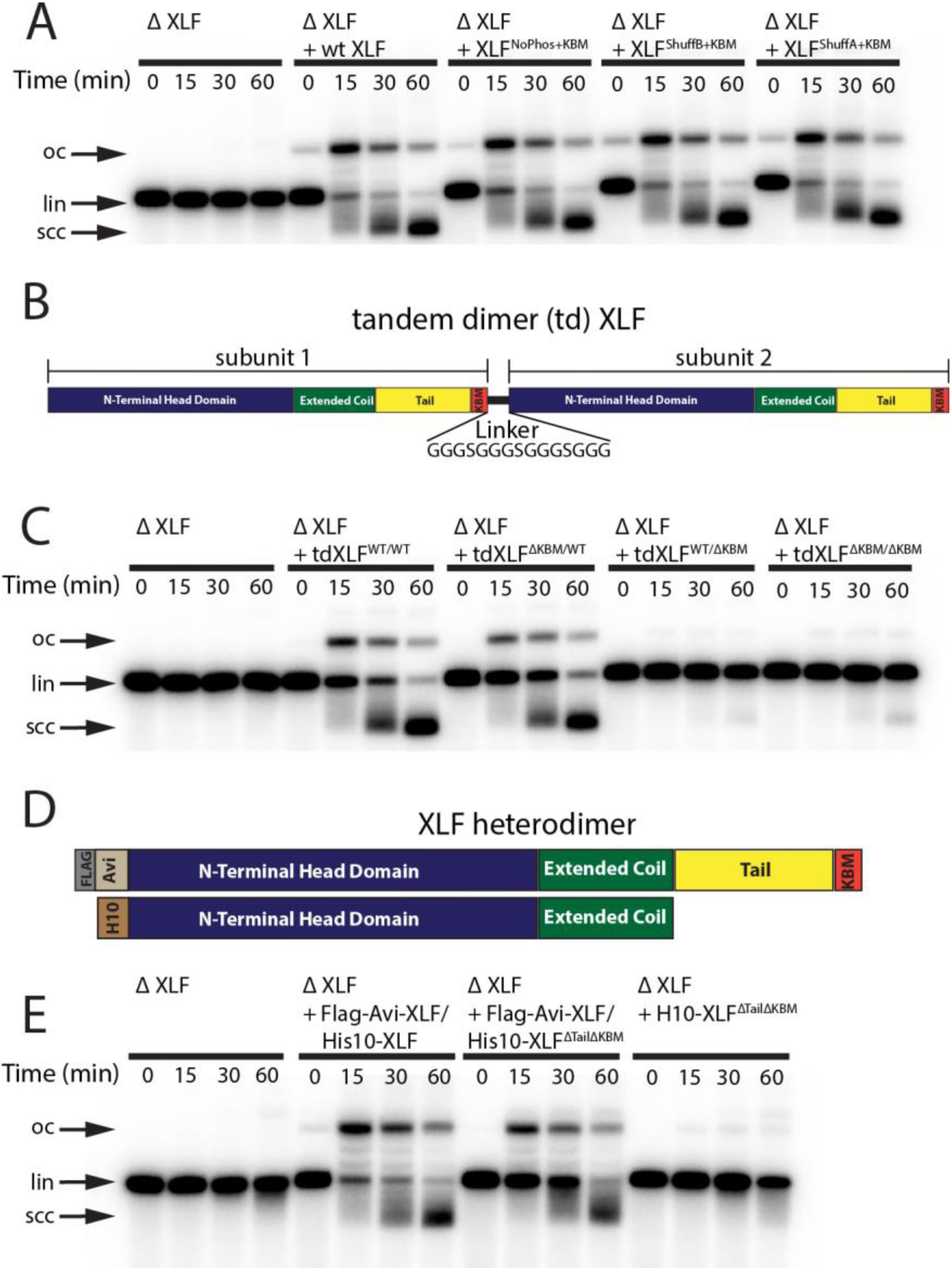
Requirements of the XLF tail in end joining. **A**, Ensemble time course end joining assay in XLF-depleted *Xenopus* egg extract. Recombinant wt XLF, XLF^NoPhos+KBM^, XLF^ShuffB+KBM^, and XLF^ShuffA+KBM^ were added back at 75 nM to their respective reaction samples. DNA species: scc, supercoiled closed circular; lin, linear; oc, open circle. **B**, Schematic of the tandem dimer (td) XLF construct. **C**, Ensemble time course end joining assay in XLF-depleted *Xenopus* egg extract. Recombinant tdXLF^WT/WT^, tdXLF^ΔKBM/WT^, tdXLF^WT/ΔKBM^, and tdXLF^ΔKBM/ΔKBM^ were added back at 50 nM final concentration to their respective reaction samples. DNA species: scc, supercoiled closed circular; lin, linear; oc, open circle. **D**, Schematics of the individual subunits coexpressed to form the XLF heterodimer, Flag-Avi-XLF/His10-XLF^ΔtailΔKBM^. **E**, Ensemble time course end joining assay in XLF-depleted *Xenopus* egg extract. Recombinant heterodimers (Flag-Avi-XLF/His10-XLF and Flag-Avi-XLF/His10-XLF^ΔtailΔKBM^) and His10-XLF^ΔtailΔKBM^ were added back at 75 nM final concentration to their respective reaction samples. DNA species: scc, supercoiled closed circular; lin, linear; oc, open circle.

### Only a single KBM and C-terminal tail is required for XLF to promote end joining in egg extract

Our results with XLF^NoPhos+KBM^ and the shuffled tail mutants suggest that it is the length and not the sequence of the tail that is important for DNA end synapsis. Having previously shown that a single XLF dimer mediates synapsis,^24^ we considered if each C-terminal tail of the XLF dimer must bridge Ku molecules on opposing DNA ends of the DSB. This network of interactions may be critical for the formation of the SR complex and therefore decreasing the length of the XLF tail or removing one of the KBMs from the XLF dimer should block end joining. To test this Ku-XLF-Ku bridge model, we constructed a tandem dimer of XLF (tdXLF), which allows for the introduction of asymmetric KBM mutations.^24^ In this construct the two subunits of XLF were expressed as a single polypeptide connected by a flexible linker (FIG 4B).

We generated mutants of the tdXLF construct where either the KBM in subunit 1 is replaced by additional flexible linker sequence (tdXLF^ΔKBM/WT^), the KBM in subunit 2 is deleted (tdXLF^WT/ΔKBM^), or where both the KBM in subunit 1 is replaced by linker sequence and the KBM in subunit 2 is deleted (tdXLF ^ΔKBM /ΔKBM^) (FIG 4B). Robust end joining was observed when tdXLF^WT/WT^ or tdXLF^ΔKBM/WT^ were used to rescue XLF-depleted egg extract, but little to no joining was observed with either tdXLF^WT/ΔKBM^ or tdXLF ^ΔKBM /ΔKBM^ (FIG 4C). The loss of end joining observed when we delete or mutate the C-terminal KBM within subunit 2, even if the KBM in subunit 1 is intact (FIG 4B-C and Supp. FIG 4A) is likely because the C-terminal residues of the XLF KBM bind Ku80 in an internal hydrophobic pocket with the – COO group at the C-terminus forming an electrostatic contact with Lys238 of Ku80 (in the human).^26^ The combination of losing this electrostatic contact and having to sterically accommodate the flexible linker within the hydrophobic pocket likely inhibits the KBM of subunit 1 from binding Ku80 effectively. A similar trend was observed when we used tdXLF constructs containing a previously characterized point mutation within the KBM (Leu 301 to Glu) (Supp. FIG 4A). The homologous mutation in human XLF was shown to reduce the affinity of XLF for Ku ∼ 5-fold *in vitro* and impaired recruitment to DNA damage in cells.^26^ Robust joining is observed when tdXLF^L301E/WT^ was used to rescue the XLF depletion, but little to no joining occured when rescuing with tdXLF^WT/L301E^ or tdXLF ^L301E /L301E^. Since we see robust joining with tdXLF^ΔKBM/WT^ and tdXLF^L301E/WT^, these results demonstrate that a single KBM within an XLF dimer is sufficient for XLF’s role in promoting end joining in extracts.

Although the above results rule out a Ku-XLF-Ku bridge mediated by both KBMs of an XLF dimer, they do not address the contributions by the tails outside of the KBM. To that end, we purified a XLF heterodimer in which one tail was deleted including the KBM. Heterodimers of XLF were generated by simultaneously expressing two versions of XLF, one His tagged and the other Flag-Avi tagged (FIG 4D).^24^ Subsequent tandem affinity purification allows for the isolation of the heterodimer as it is the only species with both affinity tags. Similar to full-length XLF heterodimers,^24^ we verified that there is little to no exchange of subunits after expression and purification (Supp. FIG 4B-C). A comparison of joining efficiency for the Flag-Avi-XLF/His10-XLF^ΔtailΔKBM^ construct and the wild type Flag-Avi-XLF/His10-XLF revealed that both constructs can rescue end joining in XLF-depleted extracts, whereas dimers of His10-XLF^ΔtailΔKBM^ cannot (FIG 4E). These results indicate a single KBM on a single tail of XLF is sufficient for robust end joining.

### The C-terminal tail of XLF must be sufficiently long to interact with XRCC4 within the synaptic complex

We next asked whether the tail is required for XLF to interact with other core NHEJ factors within the synaptic complex. Previously we showed that the XLF-XRCC4 interaction is required for SR complex formation.^24^ The flexible tail may have to be of sufficient length to facilitate the interaction between the N-terminal head domains of XLF and XRCC4. To test this hypothesis, we performed DNA pulldown experiments in egg extract to evaluate the recruitment and stability of core NHEJ factors to DNA ends. DNA containing a biotin at both 5’ ends was attached to magnetic beads and either cut to introduce a blunt-ended DSB or left intact to control for non-specific DNA binding (FIG 5A). After incubating the DNA-beads in egg extract, the beads were isolated and washed. The stably associated proteins were identified by western blot. Robust recruitment of the core NHEJ factors Ku, DNA-PKcs, XRCC4, Lig4, and XLF was observed in the mock-depleted extract relative to the uncut control DNA (FIG 5B). Consistent with previous reports, the lower electrophoretic mobility of XRCC4 and XLF is likely due to phosphorylation by DNA-PKcs.^11,28^

**Figure 5.**
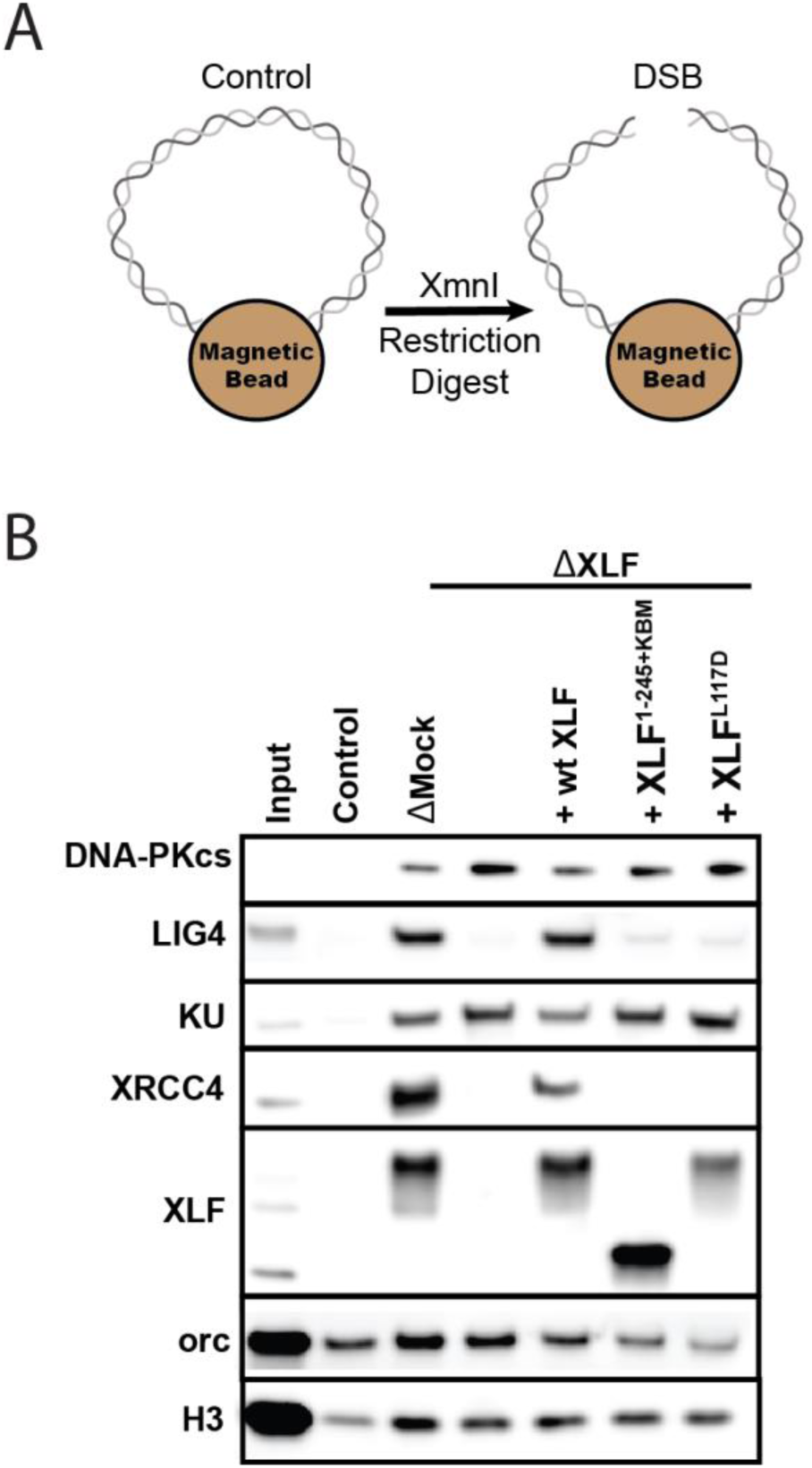
The XLF tail is required to stabilize XRCC4-Lig4. **A**, Cartoon schematic of the DNA pulldown assay. Both ends of a linear DNA substrate are conjugated to magnetic beads, and either cut with XmnI to generate a DSB with blunt ends or left uncut as a control. **B**, Immunoblots of NHEJ core factors (DNA-PKcs, Lig4, Ku, XRCC4, and XLF) and the loading controls, Orc and H3, bound to DNA-beads after a 15-minute incubation in egg extract. Samples run in parallel were the input (extract diluted 1:40) and control (uncut DNA substrate pulldown) as well as pulldowns with the DSB substrate in either mock-depleted extract or XLF-depleted extract with recombinant wt XLF, XLF^1-245+KBM^, or XLF^L117D^ added back at 20 nM.

To determine if XLF is required to stabilize XRCC4-Lig4 at DNA ends, we depleted XLF from egg extract and then blotted for NHEJ factors in our pulldown assay. Loss of XLF led to a large reduction in XRCC4 and Lig4 signal that could be rescued by addition of recombinant wild type XLF. As XRCC4-Lig4 is known to be recruited in the absence of XLF,^32^ these results are consistent with the interaction between XLF and XRCC4 stabilizing the XRCC4-Lig4 complex at DNA ends. We next tested XLF^L117D^ (human XLF^L115D^), an XRCC4 interaction deficient mutant.^24^ XLF^L117D^ was associated with DNA ends yet could not restore XRCC4-Lig4 stability (FIG 5B). Similarly, XLF^1-245+KBM^ failed to restore XRCC4-Lig4 stability even though it was robustly recruited. Collectively these results show that a minimal tail length is necessary for XLF to stabilize the XRCC4-Lig4 complex.

### NHEJ in cells also requires a single KBM and a sufficiently long C-terminal tail

Our observations in egg extract define the requirements of the XLF C-terminal tail during NHEJ. To determine if the same interactions are important in cells, we used a chromosomal end joining assay (EJ7-GFP) that reports on error-free repair of DSBs induced by Cas9 and single guide RNAs (sgRNAs).^27^ In this assay, a green fluorescent protein (GFP) expression cassette, with a 46 nt insert that disrupts GFP, is integrated into the *Pim1* locus in *Xlf* ^−/-^ mouse embryonic stem cells (mESCs). Two tandem DSBs induced by Cas9/sgRNAs excise this insert, and subsequent error-free repair between the distal DSB ends restores GFP. Thus, determining the %GFP+ cells provides the frequency of this end joining event, which is normalized to transfection efficiency (see methods). With this assay, error-free end joining depends solely on canonical NHEJ factors (Fig 6A).^27^ We transfected various human XLF constructs into *Xlf* ^−/-^ mESCs, along with the Cas9/sgRNA plasmids (Supp. FIG 5A), and measured the percentage of GFP+ cells. As previously observed, GFP+ cells are dependent on XLF in that transfections with wt XLF causes a high frequency of GFP+ cells (∼50%), but without XLF (i.e., empty vector) GFP+ cells are near background levels (FIG 6B). Similar to our results in *Xenopus* egg extract we observed that the XLF KBM and tail were both important for efficient end joining in cells (FIG 6B). As compared to wt XLF, XLF^ΔKBM^ led to a severe joining defect (6-fold), and XLF^1-243+KBM^ led to a substantial reduction (3.2-fold). Next, we introduced tdXLF constructs into *Xlf* ^−/-^ mESCs to test whether a single KBM is sufficient for XLF function. Consistent with our results in egg extract, transfection of tdXLF^(WT/WT)^ and tdXLF^(ΔKBM/WT)^ rescued repair as efficiently as wt XLF. However, tdXLF^(ΔKBM/ ΔKBM)^ showed a severe defect compared to wt XLF (12.2-fold). Overall, these results demonstrate that a single KBM within an XLF dimer is sufficient for end joining in cells, but the tail of XLF must be sufficiently long to facilitate efficient end joining.

**Figure 6.**
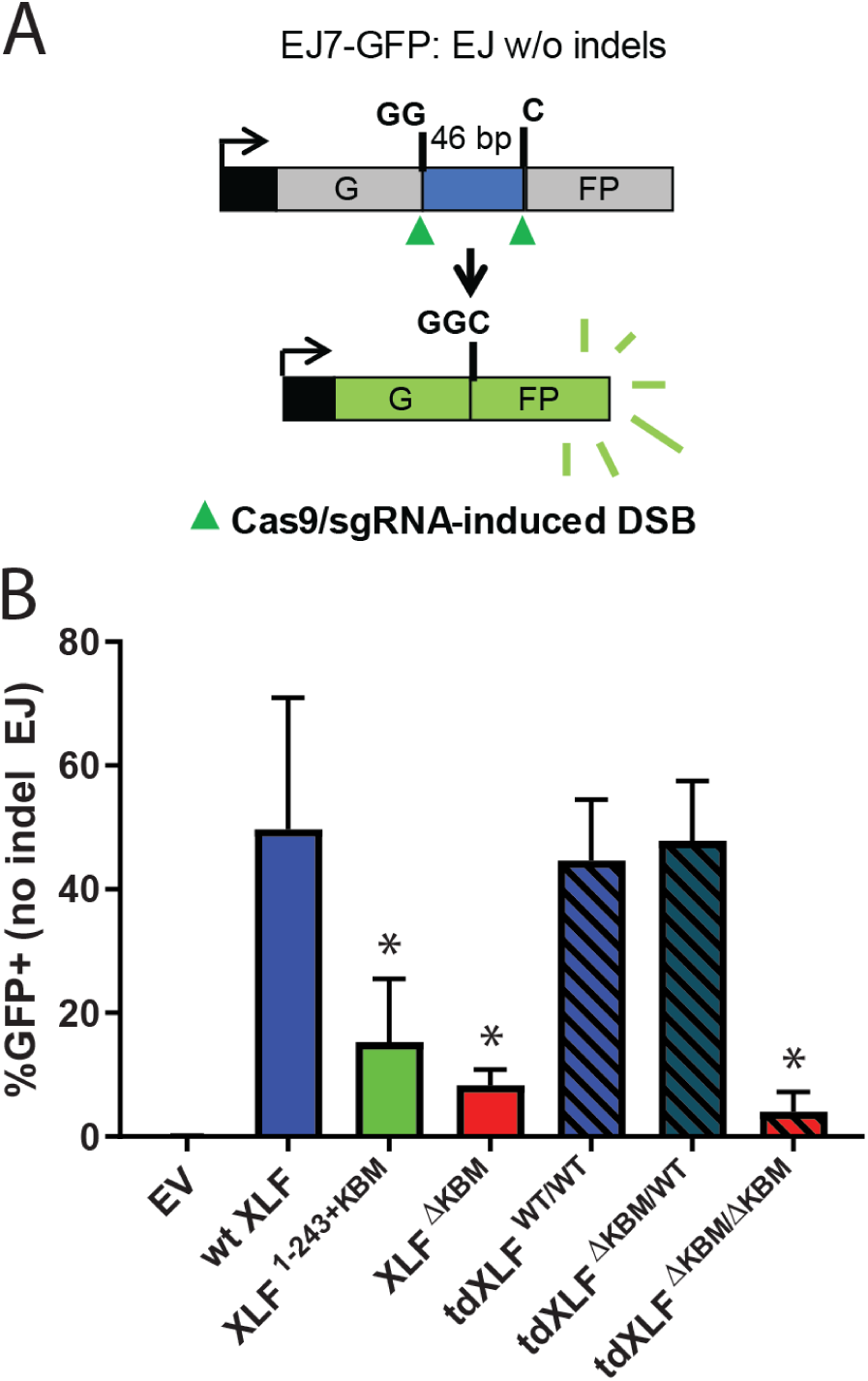
A single XLF tail is required for end joining in cells. **A**, Schematic of the cellular GFP NHEJ reporter (EJ7-GFP). A 46 bp insertion is located within a region of the GFP gene that is critical for fluorescence. Cas9 and guide RNAs are expressed so that DSBs are induced on either end of this insertion. Fluorescence is restored only if the blunt ends of the GFP gene are repaired via error-free end joining. **B**, Xlf -/- mESCs were transfected with an empty vector or the same vector containing a human XLF construct. The GFP frequencies were normalized against parallel transfections of a GFP+ expression vector. The normalized mean %GFP+ and corresponding standard deviation for each condition are shown. N=6 for each condition, and an unpaired T-Test with the Holm-Sidak correction was used to determine significance. An * represents datasets that are significantly different (*P*≤0.015) from the wt XLF relstus.

## Discussion

Synapsis of DNA ends during NHEJ is an essential but poorly understood process. Among the core NHEJ factors, the role of XLF in end joining has remained particularly elusive. Using *Xenopus* egg extract to recapitulate physiological end joining, our findings further articulate the intermolecular interactions formed between XLF and other NHEJ factors that mediate end synapsis. Here we show that both the XLF KBM and the tail are necessary for end joining. Notably, the length of the XLF tail, and not its sequence, is important for NHEJ. We propose that the C-terminal tail of a single XLF monomer acts as a “leash” that tethers XLF to Ku via its KBM and allows it to form required interactions with XRCC4 within the synaptic complex.

### The role of the XLF Ku binding motif in NHEJ

Recruitment of NHEJ factors to DSBs depends largely on Ku.^33^ XLF is strictly required for end joining in *Xenopus* egg extract.^10^ Here, we show that deleting the XLF KBM also substantially ablates end joining (FIG 1C). This defect is likely due to a loss of XLF localization to DSBs, as has been observed in cells upon KBM deletion.^25^ Therefore, the KBM recruits XLF to DNA ends which facilitates formation of the synaptic complex.

Similar to our results in egg extract, we found that introduction of XLF^ΔKBM^ into cells leads to a large decrease in error-free NHEJ, although end joining remains higher than in cells lacking XLF. As end joining could be partially rescued by high concentrations of XLF^ΔKBM^ in egg extract (FIG 1D), these results suggest that XLF does not need the KBM to promote end joining when the local XLF concentration is high enough. Altogether, these findings indicate that the XLF KBM is critical for NHEJ, although without the KBM, XLF retains a weak residual activity that is likely mediated through its interaction with XRCC4.

### Recruitment of XLF to DSBs is necessary but not sufficient for NHEJ

Recruitment of XLF to DSBs is required for end joining, but is not sufficient. In addition to the KBM, we show that the tail is critical for DNA end synapsis (FIG 3G) and subsequent end joining in egg extract (FIG 2B). Similarly, introducing an XLF mutant with a truncated tail that maintained the KBM (human XLF^1-245+KBM^) resulted in a substantial drop in NHEJ efficiency in cells (FIG 6B). Prior to this study, the role of the KBM and the rest of the C-terminal tail had not been analyzed systematically. Many structural studies of XLF have deleted both the tail and KBM^15,16,17,21,34^ as this construct facilitates crystallization.^16, 17^ This construct still interacts with XRCC4 and promotes minimal end joining in *in vitro* reconstitution assays.^12,16,34^ Within human cells, the XLF C-terminal tail was reported to be dispensable for V(D)J recombination.^35^ As we observe a substantial reduction in NHEJ efficiency in mESCs upon removal of either the tail or KBM, this result may reflect a differential requirement of intermolecular interactions during V(D)J recombination as compared to spontaneous DSB repair. Our results demonstrate that both the flexible C-terminal tail and the KBM are required for canonical NHEJ under physiological conditions.

How does the tail of XLF contribute to NHEJ? Experiments in which we shuffled the sequence of the tail demonstrated that outside the KBM there are no additional motifs that are required for end joining (FIG 4A). Similarly, ablating all known and potential phosphorylation sites within this region did not affect end joining (FIG 4A), consistent with prior results.^28,36^ These observations suggest that the length of the tail is the critical requirement for NHEJ and informs models of how XLF promotes end joining. Alternating XLF and XRCC4 filaments have been proposed to synapse DNA ends and the tail of XLF has been implicated in stabilizing these filaments in vitro.^17,34^ We disfavor this possible model as we have previously shown that only a single XLF dimer promotes SR complex formation and joining in egg extract^24^. Furthermore, recent work in a human reconstitution of NHEJ also suggest that XLF filaments are not necessary for end synapsis.^37^ Therefore, we considered a model in which the two tails of the XLF dimer enable the formation of a Ku-XLF-Ku bridge^33^ that is required to synapse DNA ends. Using synthetic tandem dimers of XLF, we generated asymmetric XLF dimers to show that a single KBM within an XLF dimer is sufficient to promote NHEJ in both *Xenopus* egg extract (FIG 4B-C) and in mammalian cells (FIG 6B). These results point towards a model where a single XLF-Ku contact is sufficient for end joining.

### A model for XLF in DNA end synapsis during NHEJ

Our data supports a sequential model that describes how XLF facilitates formation of the NHEJ synaptic complex (FIG 7). A single XLF dimer is first recruited to Ku via its KBM. Subsequently, the tail enables XLF to find and engage with other binding partners within the complex. In this way, the tail acts as a flexible connector that allows Ku-bound XLF to diffuse locally and make additional contacts that promote the transition from the LR to SR complex. One of these potential interactions is between XLF and XRCC4, an interaction that is required for NHEJ in cells.^18,19,20^ We have previously shown that a single XLF dimer must engage two XRCC4-Lig4 complexes, one through each head domain, for efficient SR complex formation and end joining.^24^ Here, we demonstrate that a sufficient XLF tail length is required for both SR complex formation and for XLF to stabilize XRCC4-Lig4 at DNA ends. These results mimic XLF L117D, an XLF mutant deficient in interaction with XRCC4.^24^ We propose that the tail of XLF facilitates the formation of a XLF-XRCC4 bridge that spans the DNA break or positions XRCC4-Lig4 so that Lig4 can engage the DNA ends. These models need not be mutually exclusive and future studies will be needed to elucidate the structural role of both XRCC4 and Lig4 in end synapsis during NHEJ.

**Figure 7.**
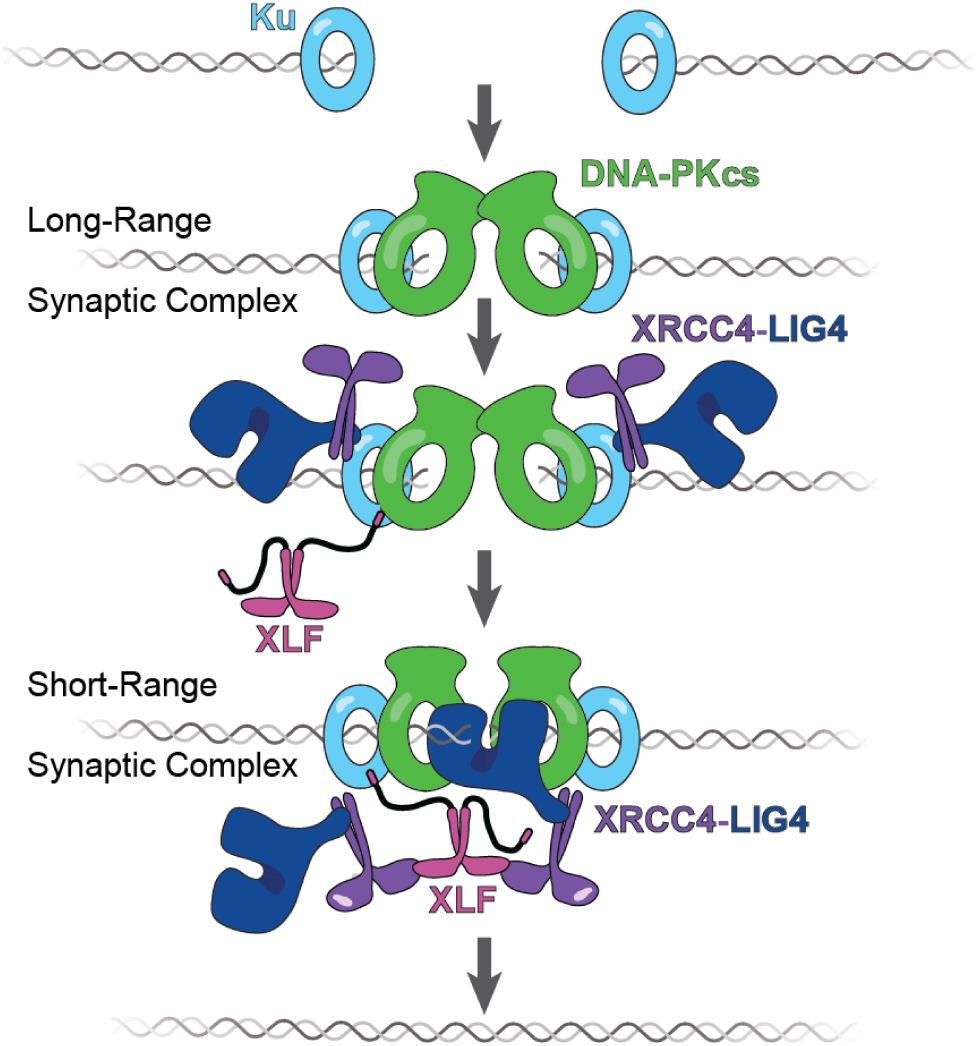
The tail enables XLF to stabilize XRCC4-Lig4 in the short-range complex. Cartoon model representing the evolution of the NHEJ synaptic complex. Ku initially binds DNA ends and DNA-PKcs is recruited shortly after to mediate formation of the LR complex. The XLF KBM tethers it to Ku while the other domains of XLF can diffuse locally to find and bind XRCC4. This mediates the formation of the XRCC4-XLF-XRCC4 bridge that leads to SR complex formation and likely puts Lig4 in position to engage the DNA ends.

## Acknowledgements

We thank T.G.W. Graham for generating and sharing the His10-SUMO-xl XLF, His10-SUMO-xl XLF^WT/WT^, and His10-xl XLF : Flag-Avi-xl XLF plasmids, K. Arnett (HMS Center for Macromolecular Interactions) for assistance with DSF assays, and members of the Loparo lab, in particular B.M. Stinson, for constructive feedback. We thank Johannes Walter and his laboratory for access to their frog facility, for sharing reagents, and for helpful discussions. This work was funded by National Institutes of Health grants (R01GM115487 to J.J.L. and R01CA197506, R01CA240392 to J.M.S.). S.M.C was funded by a National Institutes of Health Fellowship (F32GM129913). S.C.P was supported by the Molecular Biophysics Training Grant, Harvard University, National Institutes of Health (NIGMS T32 GM008313).

## Data Availability

The data and any custom analysis programs used as part of this study are available from the authors upon reasonable request.

## Author Contributions

S.M.C. and J.J.L. conceived the experiments and wrote the manuscript. S.M.C. performed all experiments with the exception of the DNA pulldown assays (performed by A.T.M.) and the cellular GFP-reporter NHEJ assays (performed by M.C-A., F.W.L., and J.M.S.). S.C.P. worked together with S.M.C. to generate and test the tandem dimer XLF constructs. All authors edited the manuscript.

## Materials and Methods

### Plasmid Construction

All expression plasmid constructs used are listed in Supp. Table 2. *xl* in the plasmid name denotes constructs based on the XLF sequence from *Xenopus laevis. h* in the plasmid name denotes constructs based on the human XLF sequence.

#### XLF and XLF Truncation Mutants

The construction of the His10-SUMO-xl XLF (*Xenopus laevis* XLF) expression plasmid, pTG296, has been previously described.^10^ This plasmid was used as the template to generate the His10-SUMO-xl XLF^ΔKBM^, His10-SUMO-xl XLF^1-245+KBM^, His10-SUMO-xl XLF^1-265+KBM^, and His10-SUMO-xl XLF^1-285+KBM^ *Xenopus laevis* XLF constructs in the same expression vector using round the horn mutagenesis.^38^ This method generates a linear PCR product where edits to the template are introduced at the termini. This linear product is then phosphorylated and ligated to produce a circular plasmid.

#### XLF NoPhos and Shuffled Tail Mutants

To generate His10-SUMO-xl XLF^NoPhos+KBM^, a geneblock that contained the sequence corresponding to the *Xenopus laevis* XLF C-terminal tail region (amino acids between and including 226 -292) was ordered (Integrated DNA Technologies) where all serine residues in this region were mutated to glycine and all threonine residues in this region were mutated to alanine. This geneblock was then inserted into the appropriate position to replace the original sequence corresponding to the C-terminal tail within the H10-SUMO-xl XLF expression plasmid by isothermal (Gibson) assembly.^39^ The His10-SUMO-xl XLF^ShuffA+KBM^ and His10-SUMO-xl XLF^ShuffA+KBM^ constructs were generated using the same approach. The shuffled sequences of the C-terminal tail regions in these mutants were generated using the Protein Sequence Shuffle Tool within the Sequence Manipulation Suite.^31^

#### tdXLF constructs

The construction of the tandem dimer of *Xenopus laevis* XLF, H10-SUMO-xl tdXLF^WT/WT^, has been previously described.^24^ This tandem dimer consists of two XLF sequences connected by a linker composed of a repeating ‘GGGS’ amino acid sequence.^24^ To create the H10-SUMO-xl tdXLF ^ΔKBM/WT^ construct, the KBM from the XLF subunit 1 sequence was deleted and additional linker sequence (GGGSGGGSGGGSGGGS) was added to prevent defects due to shortening the flexible tail and linker region between subunit 1 and subunit 2. This was accomplished by amplifying the H10-SUMO-xl tdXLF^WT/WT^ as two separate fragments and assembling them using isothermal (Gibson) assembly.^39^ The H10-SUMO-xl tdXLF ^WT/ΔKBM^ was created using the same method, but in this case the KBM of subunit 2 was deleted outright and not replaced. The double mutant, H10-SUMO-tdXLF ^ΔKBM /ΔKBM^, was also generated using this method but used H10-SUMO-xl tdXLF ^ΔKBM/WT^ as the template. The same approach was used to create expression plasmids containing H10-SUMO-xl tdXLF ^L301E/WT^, H10-SUMO-xl tdXLF ^WT/L301E^, and H10-SUMO-xl tdXLF ^L301E/L301E^. No additional linker sequence was introduced for any of the L301E point mutants.

#### XLF heterodimers

The procedure by which both His10-xl XLF and Flag-Avi-xl XLF (both *Xenopus laevis* XLF sequences) were cloned into a dual expression vector has been described previously.^24^ Round the horn mutagenesis was used to generate the Flag-Avi-xl XLF/His10-xl XLF^ΔtailΔKBM^ construct.^38^ The His10-xl XLF^ΔtailΔKBM^ construct was generated by the same method using the Flag-Avi-XLF/His10-XLF^ΔtailΔKBM^ expression plasmid as a template.

#### Human XLF and tdXLF constructs

The cloning of the human XLF constructs into a pCAGGS-BSKX vector was performed as previously described.^27^ Briefly, each XLF construct was ordered as a geneblock from Integrated DNA Technologies with a 3xFlag tag at the N-terminus. The pCAGGS-BSKX vector was linearized by cutting with EcoR1-HF (New England Biolabs) and Xho1 (New England Biolabs). Each geneblock was then inserted into the linearized vector by isothermal (Gibson) assembly.^39^ Similar to the construction of the *Xenopus* tandem dimer construct, a custom python script was used to generate distinct DNA sequences of human XLF for each XLF sequence included in the tandem dimer to facilitate cloning.^24^ These two distinct XLF sequences are separated by a ‘GGGSGGGSGGGSGGG’ linker.

### Protein Purification

All purified recombinant proteins used are show in Supp. FIG 5B-C.

#### xl XLF, XLF truncation mutants, XLF NoPhos, and Shuffled Tail Mutants

His10-SUMO-xl XLF (wild type), His10-SUMO-xl XLF^1-245+KBM^, His10-SUMO-xl XLF^1-265+KBM^, His10-SUMO-xl XLF^1-285+KBM^, His10-SUMO-xl XLF^NoPhos+KBM^, His10-SUMO-xl XLF^ShuffA+KBM^, and His10-SUMO-xl XLF^ShuffB+KBM^ constructs were all purified using a previously detailed protocol.^24^ Each expression plasmid was transformed into *E*.*coli* BL21(DE3)pLysS cells. Cultures were grown at 37°C to an OD_600_ of ∼0.6. IPTG was then added to cultures at a final concentration of 1mM. The cultures were then moved to 25-30°C for 3 hours for protein expression. Cultures were then centrifuged to collect cells. Cell pellets were washed in 1x PBS buffer, flash frozen in liquid nitrogen, and stored at -80°C. The pellets were thawed and resuspended in 15mL lysis buffer (20 mM Tris-HCl, pH 8.0, 1 M NaCl, 30 mM imidazole, 5 mM BME, and 1mM PMSF) per liter of culture and sonicated. The resulting lysates were clarified by centrifugation for 1 hour at 20000 rpm in a SS34 fixed angle rotor at 4°C. The supernatant was incubated with Ni-NTA agarose (Qiagen) equilibrated in lysis buffer for 1 hour at 4°C to allow for binding of His-tagged proteins. The Ni-NTA resin was then was washed with lysis buffer followed by washing with salt reduction buffer (20 mM Tris-HCl, pH 8.0, 350 mM NaCl, 30 mM imidazole, 5 mM BME). Proteins bound to the Ni-NTA resin were then eluted by incubating the Ni-NTA resin in elution buffer (20 mM Tris-HCl, pH 8.0, 350 mM NaCl, 250 mM imidazole, 5 mM BME) for 2 minutes. This elution step was repeated several times. Peak fractions were pooled and dialyzed against His-SUMO dialysis buffer(20 mM Tris-HCl, pH 8.0, 350 mM NaCl, 10mM imidazole, 5mM BME, and 10% glycerol) at 4°C in the presence of H6-Ulp1 protease, which was added to cleave the H10-SUMO tags from the XLF constructs. After two rounds of dialysis, each being more than 4 hours long, the dialysate was incubated with fresh Ni-NTA resin equilibrated in dialysis buffer at 4°C for 1.5 hours. Any remaining H10-SUMO-protein, cleaved H10-SUMO, or H6-Ulp1 should remain bound to the Ni-NTA at this step. The flow through which contains XLF protein lacking the H10-SUMO tag was collected and diluted in 1.33 volumes of SP Buffer (50 mM Na-MES, pH 6.5, 10% glycerol, 1 mM DTT) so that the [NaCl] becomes 150 mM, and subsequently passed over SP Sepharose Fast Flow (Amersham) equilibrated in SP Buffer A (50 mM Na-MES, pH 6.5, 150 mM NaCl, 10% glycerol, 1 mM DTT). The SP Sepharose Fast Flow resin was then washed with 10 column volumes of SP Buffer A. The protein was eluted with SP Buffer B (50 mM Na-MES, pH 6.5, 350 mM NaCl, 10% glycerol, 1 mM DTT) in 1 column volume increments. All steps involving SP Sepharose Fast Flow were carried out at 4°C. Peak fractions were pooled, flash frozen in liquid nitrogen, and stored at -80°C until use.

The purification of His10-SUMO-xl XLF^ΔKBM^ followed the His10-SUMO purification protocol detailed above through the SUMO cleavage and dialysis step. The dialysate was spun at 4°C for 45 minutes at ∼20,000 g and subsequently diluted ∼3-fold in Q Buffer (20 mM Tris, pH 8.0, 10% glycerol, 5 mM BME) so that the salt concentration of resulting sample was ∼100 mM NaCl. This sample was then loaded onto a HiTrap Q HP column that was equilibrated in Q Wash Buffer (20 mM Tris, pH 8.0, 100 mM NaCl, 10% glycerol, 5 mM BME) and washed with 5 column volumes of Q Wash Buffer. The xl XLF^ΔKBM^ protein was collected in the flow through which was then loaded onto a HiTrap HP SP column where majority of the sample was collected in the flow through again and subsequently concentrated using a 3-MWCO centrifugal spin filter (Amicon). The sample was then flash frozen in liquid nitrogen and stored at -80°C until use.

#### xl tdXLF constructs

H10-SUMO-xl tdXLF^WT/WT^, H10-SUMO-xl tdXLF ^ΔKBM/WT^, H10-SUMO-xl tdXLF ^WT/ΔKBM^, H10-SUMO-xl tdXLF ^ΔKBM/ ΔKBM^, H10-SUMO-xl tdXLF ^L301E/WT^, H10-SUMO-xl tdXLF ^WT/L301E^, and H10-SUMO-xl tdXLF ^L301E/L301E^ were all expressed and purified as previously described.^24^ Each tdXLF construct was transformed into *E*.*coli* BL21(DE3)pLysS cells and cultures were grown at 37°C until the OD_600_ was between 0.55 and 0.70. Expression was induced by adding IPTG to a 1 mM final concentration. Expression was the carried out at 22°C for 4 hours. Cultures were then spun down, washed in 1x PBS buffer, flash frozen in liquid nitrogen, and stored at -80°C. The same His10-SUMO purification steps described for wild type XLF above were then followed for the tandem dimer constructs. The flow through from the second Ni-NTA agarose resin (Qiagen) incubation was spun at 20,000 x g for 1 hour at 4°C. The sample was then diluted with 2.5 volumes of SP Buffer (50 mM Na-MES, pH 6.5, 10% glycerol, 5 mM BME) so that final [NaCl] was 100 mM in the sample before being loaded onto a HiTrap SP HP column that was equilibrated in SP Buffer A (50 mM Na-MES, pH 6.5, 10% glycerol, 5 mM BME, 100 mM NaCl). The column was then washed with 5 column volumes of SP Buffer A. Protein bound to the column was eluted using a 100 – 1000 mM NaCl gradient over 30 column volumes. Peak fractions were pooled and concentrated using a 3- or 10-kDa MWCO centrifugal spin filter (Amicon). In the case of the H10-SUMO-tdXLF ^ΔKBM/ ΔKBM^ mutant, the isoelectric point is significantly lower than that of the wild type XLF (6.13 vs. 8.00), and this mutant did not stick to the SP HP column. For this mutant, the SP HP flow through was concentrated and taken to the next step. The concentrated sample from the HiTrap SP HP column was then loaded onto a Superdex 200 Increase 10/300 equilibrated in (50 mM Na-MES, pH 6.5, 10% glycerol, 350mM NaCl, 5 mM BME). Peak fractions were pooled and concentrated as described above. Samples were flash frozen in liquid nitrogen and stored at -80°C until use.

#### xl XLF heterodimers

Purification of XLF heterodimers followed a previously described protocol.^24^ The Flag-Avi-xl XLF/His10-XLF and Flag-Avi-xl XLF/His10-xl XLF^ΔtailΔKBM^ XLF heterodimer constructs were transformed into BL21(DE3) pLysS cells along with a BirA biotin ligase expression plasmid. Cultures were grown at 37°C until the OD_600_ was between 0.4 and 0.6. IPTG was then added to cultures at a 1 mM final concentration, and biotin was added at a final concentration of 25 µM to allow for BirA-dependent biotinylation of Avi-tagged proteins. Cultures were moved to 22°C for 4 hours. Cultures were then spun down to cell pellets, washed in 1x PBS buffer, flash frozen in liquid nitrogen, and stored at -80°C. Pellets were thawed and resuspended in 15 mL of His-SUMO lysis buffer (20 mM Tris-HCl, pH 8.0, 1 M NaCl, 30 mM imidazole, 5 mM BME, and 1mM PMSF) per liter of culture and sonicated. The lysates were then spun at 20000 rpm in a SS34 fixed angle rotor for 1 hour at 4°C. For 90 minutes at 4°C, the supernatant was incubated with Ni-NTA agarose (Qiagen) that was equilibrated in His-SUMO lysis buffer. The Ni-NTA resin was then washed with lysis buffer. Proteins bound to the Ni-NTA resin were then eluted by incubating the Ni-NTA resin in elution buffer (20 mM Tris-HCl, pH 8.0, 350 mM NaCl, 250 mM imidazole, 5 mM BME) and incubating for 2 minutes. This elution step was repeated several times. The peak fractions from the Ni-NTA eluate were then pooled and passed over SoftLink Avidin resin (Promega). The SoftLink Avidin resin was then washed extensively with SoftLink Avidin Wash Buffer A (20 mM Tris HCl, pH 8, 1 M NaCl, 10% glycerol, 5 mM BME) to remove any proteins that do not have a biotinylated AviTag. SoftLink Avidin Wash Buffer B (20 mM Tris-HCl, pH 8, 350 mM NaCl, 10% glycerol, 5 mM BME) was then put over the resin to bring the [NaCl] down to 350 mM. Avi-tagged protein was eluted from the SoftLink Avidin resin using SoftLink Avidin Elution Buffer (20 mM Tris HCl, pH 8, 350 mM NaCl, 10% glycerol, 5 mM BME, 5 mM biotin). Peak elution fractions were pooled and concentrated using a 10 kDa MWCO centrifugal concentrator (Amicon) before being flash frozen in liquid nitrogen and stored at -80°C.

#### His10-xl XLF^ΔtailΔKBM^

Expression and purification of His10-xl XLF^ΔtailΔKBM^ followed a previously described protocol.^24^ The same expression and purification steps used for the wild type His10-SUMO-xl XLF construct were used for this construct. The Ni-NTA eluate was dialyzed against MonoQ Buffer A (20mM Tris-HCl, pH 8.0, 10% glycerol, 100 mM NaCl, 5 mM BME) at 4°C. After 2 rounds of dialysis, each being more than 6 hours long, the dialysate was filtered using a 0.22 µm syringe filter. The filtered sample was then loaded onto HiTrap Mono Q column equilibrated in MonoQ Buffer A. The column was washed with seven column volumes of Buffer A, and protein was eluted from the column using a 30 mL gradient of Buffer A into Buffer B (20mM Tris-HCl, pH 8.0, 10% glycerol, 1000 mM NaCl, 5 mM BME). Peak fractions were pooled and concentrated using a 3 kDa MWCO centrifugal concentrator (Amicon). The sample was then flash frozen in liquid nitrogen and stored at -80°C until use.

### XLF Subunit Exchange Assay

Ensemble end joining assays utilizing XLF heterodimer constructs require that the subunits do not exchange over the course of the experiment. We have previously shown that there is no subunit exchange for full length XLF constructs over a timescale of hours. We employ a similar protocol here to test whether individual XLF monomeric subunits can exchange between full length and C-terminally truncated XLF dimers.^24^ His10-XLF^ΔtailΔKBM^ and wt XLF were purified separately as described above. These proteins were then mixed in a 10 µL volume of protein storage buffer (20 mM Tris-HCl, pH 8, 350 mM NaCl, 10% glycerol, and 5 mM BME) so that the final concentration of each was 5 µM. This mixture was left to incubate at room temperature for 1-3 hours in a humidified chamber to prevent evaporation. After incubation the sample was spun at 16000 rcf for 10 minutes at room temperature or 4°C. 100 µL of His-SUMO Lysis buffer (20 mM Tris-HCl, pH 8.0, 1 mM NaCl, 30 mM imidazole, and 5 mM BME) was added to the 10 µL mixture. A 20 µL aliquot was taken at this point and mixed with 20 µL of 2x Laemmli sample buffer (Bio-Rad). 80 µL of the mixture was then incubated for 45-60 minutes at room temperature or 4°C with 10 µL NiNTA resin (Qiagen) prewashed with His-SUMO Lysis buffer (20 mM Tris-HCl, pH 8.0, 1 M NaCl, 30 mM imidazole, 5 mM BME). This sample was then spun down, and the supernatant was collected. The NiNTA resin was then washed by resuspending the resin in 500 µL of His-SUMO Lysis buffer, spinning, removing the supernatant and mixing 20 µL with 20 µL of 2x Laemmli sample buffer, and repeating twice. 80 µL of His-SUMO Elution Buffer (20 mM Tris-HCl, pH 8.0, 350 mM NaCl, 300 mM imidazole, 5 mM BME) was then added to resuspend the NiNTA resin and incubated for 10-30 minutes at room temperature. Next, the sample was spun again, and the supernatant from this spin (eluate) was collected. 20 µL of the eluate was then mixed with 20 µL of 2x Laemmli sample buffer. The resin was then resuspended in 80 µL His-SUMO Elution Buffer and a 20 µL aliquot was taken and mixed with 20 µL of 2x Laemmli sample buffer. All samples were heated to 95°C for 5 minutes and cooled to room temperature.

Samples were then run on a 4-15% precast SDS-PAGE gel (Bio-Rad), transferred to polyvinylidene fluoride membranes for 16.5 hours at 30 V at 4°C, and blocked with 5% powdered nonfat milk in PBST buffer (1x phosphate-buffered saline with 0.05% Tween 20). Membranes were probed for 1 hour at room temperature with 1:500 anti-XLF(New England Peptides, details in Methods under Immunodepletion) or 1:1000 anti-His (Bio-Rad MCA1396A) in PBST with 2.5% BSA. Membranes were then washed 3x with PBST. The anti-His blot was then probed with 1:20,000 horseradish peroxidase-conjugated rabbit anti-mouse IgG (H+L) secondary antibody (Jackson ImmunoResearch) in 5% non-fat milk in PBST for 1 hour at room temperature. The anti-XLF blot was also probed for 1 hour at room temperature with 1:10,000 goat anti-rabbit IgG horseradish peroxidase-conjugated secondary antibody (Jackson ImmunoResearch) in 5% non-fat milk in PBST. Anti-XLF can be used to exclusively monitor wt XLF because His10-XLF^ΔtailΔKBM^ does not contain the C-terminal peptide used to generate the antibody (see in Methods under Immunodepletion). Membranes were then washed extensively in PBST. The anti-His membrane was incubated in substrate solution (Pierce ECL Western Blotting Substrate Kit #32106) for ∼120 seconds. The anti-XLF membrane was incubated for ∼120 seconds with either HyGLO chemiluminescent HRP antibody detection reagent (Denville) or substrate solution (Pierce ECL Western Blotting Substrate Kit #32106). Both membranes were imaged using an Amersham Imager 600 (GE Healthcare).

### Differential Scanning Fluorimetry

Differential scanning fluorimetry protein thermal shift assays were carried out using a QuantStudio 7 Flex Real-Time PCR System (Applied Biosystems). Reactions were pipetted into wells in a MicroAmp FAST optical 96-well plate (Life Technologies) and covered with MicroAmp Optical Adhesive Film (Life Technologies). Each 30 µL reaction mixture containing the XLF construct of interest at a 2.5 µM concentration and the SPYRO Orange dye at a 1x concentration (see Protein Thermal Shift Dye Kit, Applied Biosystems). Protein samples were diluted in XLF storage buffer (20mM Tris pH 8.0, 350 mM NaCl, 5 mM BME, and 10% glycerol). After a 2-minute incubation at 25°C, the temperature was raised to 99°C at 0.05°C per second. The fluorescent dye was excited and measured using 470 nm and 587 nm, respectively. Melting temperatures were determined by fitting the emission signals with the Boltzmann equation using the Protein Thermal Shift software (Life Technologies). Each replicate consisted of three 30 µL reactions for each distinct sample. The average of two replicates is plotted for each sample, with error bars representing the min and max values from the two replicates. A two-tailed, unpaired *t* test with unequal variance and the Bonferroni correction was performed to determine if the melting temperature of the XLF mutants were significantly different from wt XLF.

### *Xenopus* egg extract preparation

Cell free extract was prepared from eggs of *Xenopus laevis* as previously described.^40^ The Center for Animal Resources and Comparative Medicine at Harvard Medical School (AAALAC accredited) cared for the female frogs used to produce eggs for this study. All work performed in this study was done in accordance AAALAC rules and regulations and approved by the Institutional Animal Care and Use Committee (IACUC) of Harvard Medical School.

### Immunodepletion

The XLF antibody used here is the same as previously described.^10^ This peptide antibody was generated by New England Peptide, Inc (Gardner, MA) using a peptide (Ac-CGASKPKKKAKGLFM-OH) corresponding to the C-terminal sequence of XLF. Immunodepletion of endogenous XLF within egg extract was carried out as detailed previously.^10^ Nocodozole was added to extract at 7.5 ng/µL prior to immunodepletion or prior to use in experiments if no immunodepletion was required. Unless otherwise noted, all rescue experiments here used recombinant XLF added back to extract at 75 nM (monomer concentration) to match a previous measurement of XLF concentration in *Xenopus laevis* eggs.^29^ Mock depletions were carried out using the same protocol and IgG purified from Rabbit Serum (Gibco) by protein A sepharose affinity chromatography as previously described.^10^

### Ensemble End Joining Assay

The ensemble gel-based end joining time course and titration assays were performed as previously described.^10,24^ For each reaction condition, extract was supplemented with a 30x ATP regeneration mixture (65 mM ATP, 650 mM phosphocreatine, 160 ng/µL creatine phosphokinase) to a 1x final concentration and 25-30 ng/µL closed circular “carrier” DNA that is required for joining of dilute linear substrates as well as for DNA replication in extract.^10^,^41^ In cases where recombinant protein is added to extract and directly compared to conditions where protein was not added, the corresponding protein storage buffer was added back to those conditions without recombinant protein to ensure the volume and composition of each reaction are directly comparable. To initiate the time course end joining reactions, a radiolabeled 2.8 kb linear DNA substrate with blunt ends was added to reactions at approximately 1 ng/µL. The preparation of this substrate has been previously described.^10^,^30^ Reactions were carried out at room temperature for the indicated time. Time points were taken by removing an aliquot of the reaction mixture and stopping the reaction by addition of stop solution (80 mM Tris, pH 8.0, 8 mM EDTA, 0.13% phosphoric acid, 10% Ficoll, 5% SDS, and 0.2% bromophenol blue). The 0-minute time point was taken and mixed with stop solution immediately after adding the radiolabeled substrate to the reaction and mixing. The titration end joining assay was assembled on a thermocycler at 2°C. These reactions were initiated by moving the thermocycler to 22°C for 20 minutes. At the 20-minute time point, the thermocycler was moved back to 2°C, and stop solution was added to each reaction. All reaction samples were then digested for 1 hour at 37°C by adding proteinase K. Digested samples were then run on a Tris-borate-EDTA 0.8% agarose gel. The gel was then pressed and dried onto a HyBond-XL nylon membrane (GE Healthcare) and exposed to a storage phosphor screen. Exposed screens were scanned using a Typhoon FLA 7000 imager (GE Healthcare).

### Microscope and Flow Cell Construction

Single-molecule experiments were performed using a through-objective TIRF microscope built around an Olympus IX-71 inverted microscope. 532 nm and 641 nm laser beams (Coherent Sapphire 532 and Coherent Cube 641) were expanded, combined using dichroic mirrors, expanded again, and focused on the rear focal plane of an oil immersion objective (Olympus UPlanSApo, 100x; NA, 1.40). The focusing lens was placed on a vertical translation stage to permit manual adjustment of the TIRF angle. The emission light was separated from the excitation light using a multipass dichroic mirror. The laser lines were further attenuated with a StopLine 488/532/635 notch filter (Semrock). A home-built beamsplitter^30^ was used to separate Cy3 and Cy5 emission signals. These two channels were imaged on separate halves of an electron-multiplying charge-coupled device camera (Hamamatsu, ImageEM 9100-13), which was operated at maximum EM gain. An automatic microstage (Mad City Labs) was used to position the sample and move between fields of view.

Microfluidic flow cells were constructed as previously described.^7,30^ Briefly, holes for an inlet and an outlet were drilled in a glass microscope slide and PE tubing was placed into each and sealed with epoxy. A channel was cut out of a strip of double-sided SecureSeal Adhesive Sheet (Grace Bio-Labs), and this channel was placed onto the glass slide so that the two holes are at opposing ends of the channel. A glass coverslip was placed on the bottom of the flow cell on the second side of the double-sided adhesive. This coverslip was functionalized with a mixture of mPEG-SVA-5000 (Laysan Bio, Inc.) and biotin-mPEG-SVA-5000 (Laysan Bio, Inc.). The edges of the channel were then sealed with epoxy.

### Single-Molecule DNA SR Complex Formation Assay

The protocol for the intramolecular end joining single-molecule FRET assay generally follows a previously established protocol.^10,30^ 1 mg/mL streptavidin in PBS buffer was flowed into the flow cell and left to incubate at room temperature for 5 minutes. This solution was then washed out with egg lysis buffer (ELB; 10 mM HEPES, pH 7.7, 50 mM KCl, 2.5 mM MgCl_2_) before introducing the DNA substrate. This 2kb blunt-ended linear DNA substrate contains an internal biotin for immobilization within the flow cell. Cy3 and Cy5 flourophores are positioned 7 nucleotides from each opposing end so that energy can be transferred from Cy3 to Cy5 upon Cy3 excitation if those opposing ends are synapsed. The construction of this substrate has been described in detail previously.^10,30^ The DNA substrate was flowed into the flow cell in the presence of an oxygen scavenging system (5 mM protocatechuic acid (PCA) and 100 nM protocatechuate 3,4-dioxygensae (PCD)). 1 mM trolox was also included and serves as a triplet state quencher. This oxygen scavenging system with trolox was included in every solution introduced into the flow cell and imaged. After ∼5 minutes the DNA solution was washed out of the flow cell with ELB supplemented with the oxygen scavenging system.

Samples of extract were immunodepleted, supplemented with the oxygen scavenging system with trolox, the ATP-regeneration system described above, and recombinant protein as described above. Recombinant protein was included at 500 nM. Data acquisition begun 30 to 60 seconds after flowing the extract sample into the flow cell. 1-second exposures were collected every other second, and laser excitation was alternated so that during a single excitation cycle four 532 nm excitation frames were followed by a single 641 nm excitation frame. These excitation cycles were repeated while imaging a single field of view within the flow cell for 15 minutes. After 15 minutes, the flow cell was moved via the automatic microstage so that a new and distinct field of view was now centered under the objective and brought into focus. The same imaging procedure was repeated for 15 additional minutes in this new field of view. This process was repeated so that distinct fields of view were imaged from 0-15 minutes, 15-30 minutes, and 30-45 minutes. Previously described custom MATLAB scripts were used to analyze the data and are available upon request.^10,24.30^ Trajectories were truncated prior to photobleaching events that were detected automatically during data processing and analysis. Additionally, trajectories were excluded if they exhibited low signal-to-noise, multistep photobleaching, large fluctuations in fluorescence intensity not due to FRET, or if there was more than one peak present within a region of interest. SR complex formation events were identified manually using 5 consecutive frames above a FRET threshold (0.25) as a guideline. The rate of SR complex formation was calculated by dividing the number of SR complex formation events by the total time that SR complex formation was possible (the total time that Cy3 and Cy5 emission signal are both present and the ends were not already joined, summed over all substrate molecules that were tracked for a certain experimental condition). Sample sizes are reported in Supplementary Table 1. The plot in Figure 3G was generated using the notBoxPlot MATLAB function.^42^

### DNA Pulldown Assay

The DNA pulldown assay was performed essentially as previously described.^7^ This protocol is outlined below with any alterations described in detail.

Biotinylated primers were used to generate a 1kb DNA fragment with biotin molecules attached to the 5’ termini. Streptavidin-coated magnetic beads (36 µL per biological replicate) were washed twice in 2x Bead Wash Buffer (10 mM Tris, pH 7.4, 2 M NaCl, and 20 mM EDTA), and subsequently resuspended in 1x Bead Wash Buffer with 30 nM of the biotinylated DNA substrate described above at room temperature for 20 minutes. The DNA-bound beads were again washed twice in 2x Bead Wash Buffer and then washed twice in 1x Cutsmart Buffer (NEB) before being resuspended in 80 µL of 1x Cutsmart Buffer. The DNA-bead suspension was then split so that plasmid control and DSB samples could be prepared and tested in parallel. 1 µL of Xmn1 (NEB) was added to the DSB sample to generate the DSB, and both samples were incubated at 37C for 6 hours. Samples were then washed 3x with 2x Bead Wash Buffer and once in Egg Lysis Blocking Buffer (10mM HEPES, pH 7.7, 50 mM KCl, 2.5 mM MgCl_2_, 250 mM sucrose, and 0.02% Tween20) to remove the Xmn1 and reduce non-specific binding to the beads. Beads were then incubated in Egg Lysis Blocking Buffer for 20 minutes on ice and then resuspended in 50 µL of Egg Lysis Blocking Buffer.

Extract was supplemented with the 30x ATP regeneration mixture (65 mM ATP, 650 mM phosphocreatine, 160 ng/µL creatine phosphokinase) to a 1x final concentration and with 30 ng/µL of circular pBlueScript II plasmid to act as carrier DNA.^41^ 15 µL of the DNA-beads sample was mixed with an equal volume of extract and incubated for 15 minutes at room temperature. 30 µL of the reaction was then layered over 180 µL of ELB-Sucrose Cushion Solution (10 mM HEPES, pH 7.7, 50 mM KCl, 2.5 mM MgCl_2_, 500 mM sucrose) in a horizontal rotor (Komp-spin, Ku Prima-18R). The pelleted DNA-beads were then washed in 180 µL of Egg Lysis Blocking Buffer and resuspended in 20 µL of 1x reducing Laemmli sample buffer. Extract was diluted 1:40 in 1x reducing Laemmli sample buffer to be used as input sample for western blotting. Samples were separated on a 4-15% precast SDS-PAGE gel (BioRad) for 30 minutes at 200V and subsequently transferred to a PVDF membrane for 60 minutes at 103V at 4°C. Membranes were blocked with 5% powdered nonfat milk dissolved in 1x PBST for 30 minutes and then incubated with primary antibody diluted in 1xPBST containing 2.5% BSA (OmniPur) for 12-16 hours at 4°C. Primary antibodies were used at the following concentrations: ∝-Ku80 1:10000, ∝-Lig4 1:2000, ∝-XRCC4 1:2000, ∝-XLF 1:500, ∝-DNA-PKcs 1:5000, and ∝-Orc2 1:10000. After extensive washing with 1x PBST, membranes were incubated with goat anti-rabbit-HRP (Jackson ImmunoResearch) secondary antibody diluted 1:10,000 or 1:20,000 or rabbit anti-mouse-HRP (Jackson ImmunoResearch) secondary antibody diluted 1:10,000 in 5% powered nonfat milk and 1x PBST for 1 hour at room temperature. Membranes were washed again with 1x PBST, incubated with HyGLO Quick Spray (Denville Scientific), and imaged on an Amersham Imager 600 (GE Healthcare).

### Cellular GFP Reporter NHEJ Assay

Xlf-/-mESCs with the EJ7-GFP reporter integrated at the Pim1 locus, Cas9/sgRNA plasmids for targeting the DSBs to this reporter (i.e., px330-7a and px330-7b), and pCAGGS-BSKX empty vector (EV), were previously described.^27^ Cas9/sgRNA plasmids used the px330 plasmid (Addgene #42230).

For the reporter assays, cells were seeded on a 24 well plate, and subsequently transfected with 200 ng of px330-7a, 200 ng px330-7b, and 50 ng of EV or XLF expression vector, using 1.8 uL of Lipofectamine 2000 (Thermofisher) in 0.6 mL total volume. Three days after transfection, cells were analyzed by flow cytometry using a CyAN ADP (Dako), as described (Bhargava 2018). GFP+ frequencies were normalized to parallel transfections with a GFP+ expression vector (pCAGGS-NZE-GFP) (Bhargava 2018).

#### Immunobloting Analysis

For immunoblotting analysis, cells were transfected as for the reporter assays, but scaled twofold onto a 12 well, and replacing the Cas9/sgRNA plasmids with EV. Subsequently, cells were extracted using NETN buffer (20 mM Tris at pH 8.0, 100 mM NaCl, 1 mM EDTA, 0.5% Igepal, 1.25 mM DTT, Roche protease inhibitor) with several freeze/thaw cycles. Extracts were probed with antibodies for mouse monoclonal anti-FLAG HRP (Sigma-Aldrich Cat#A8592), or rabbit polyclonal anti-ACTIN (Sigma-Aldrich Cat#A2066) with the secondary antibody goat polyclonal Anti-Rabbit IgG HRP (Abcam Cat#ab205718). ECL western blotting substrate (Thermo Fisher Cat#32106) was used to develop HRP signals.

**Supplementary Figure 1.**
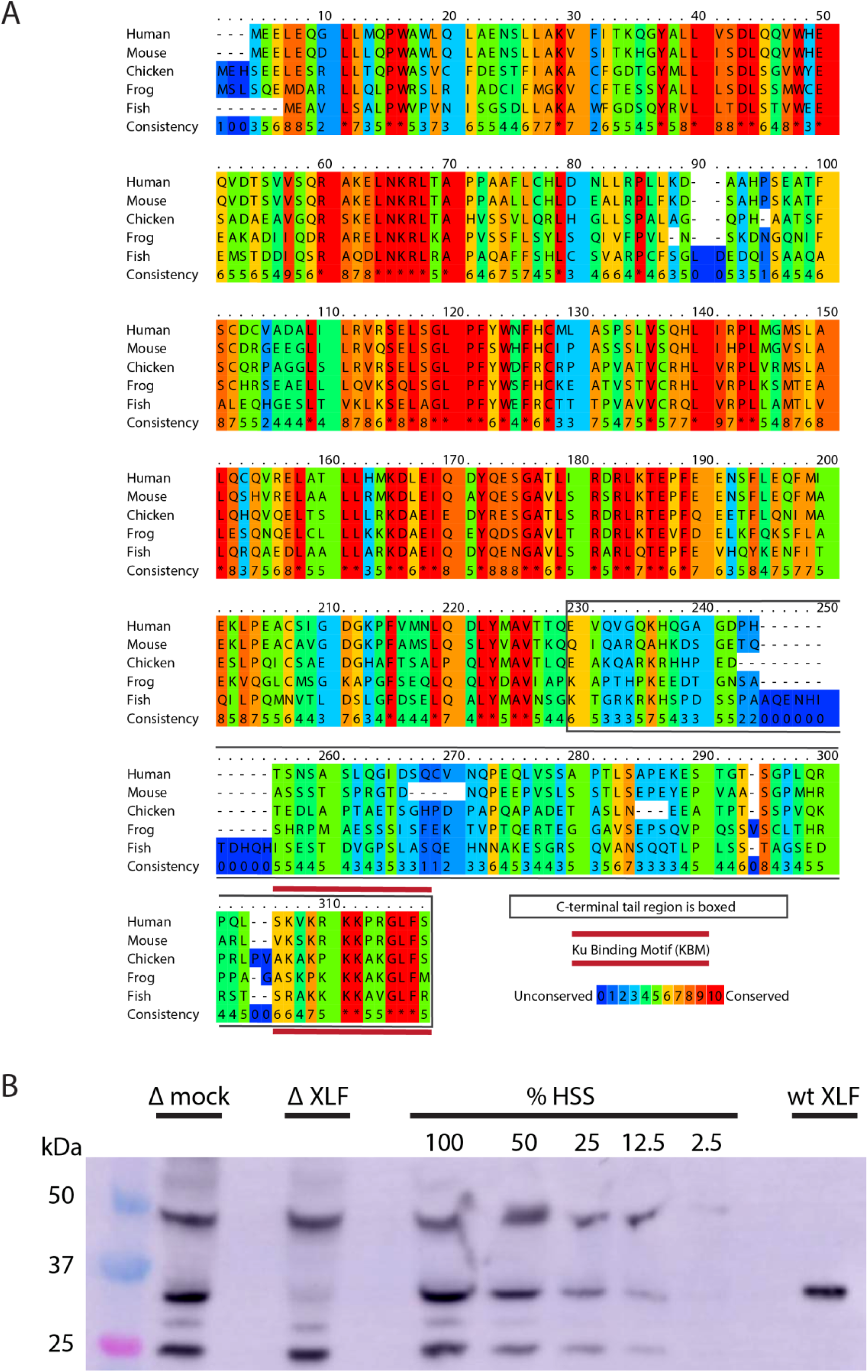
*Xenopus laevis* XLF is evolutionarily conserved and can be depleted from egg extract. **A**, A protein sequence alignment of XLF from human (*Homo sapiens* UniProt ID Q9H9Q4), mice (*Mus musculus* UniProt ID Q3KNJ2), chicken (*Gallus gallus* UniProt ID F1NVP8), frog (*Xenopus laevis* see note), and fish (*Danio rerio* UniProt ID Q6NV18). The *Xenopus laevis* sequence and translation start site was determined previously by immunoprecipitation from extract and subsequent trypsin digestion and analysis by mass spectrometry.^10^ The alignment was performed using PRALINE multiple sequence alignment and its default settings.^43^ A colored scale shows the degree of conservation. The C-terminal tail is boxed in grey, and the KBM is highlighted by a red bar above and below the sequences. **B**, Western blotting for XLF showing its presence in extract and its relative absence in XLF-depleted extract. We also show that our purified recombinant wt XLF (87nM here) runs at the same molecular weight as the endogenous XLF from extract.

**Supplementary Figure 2.**
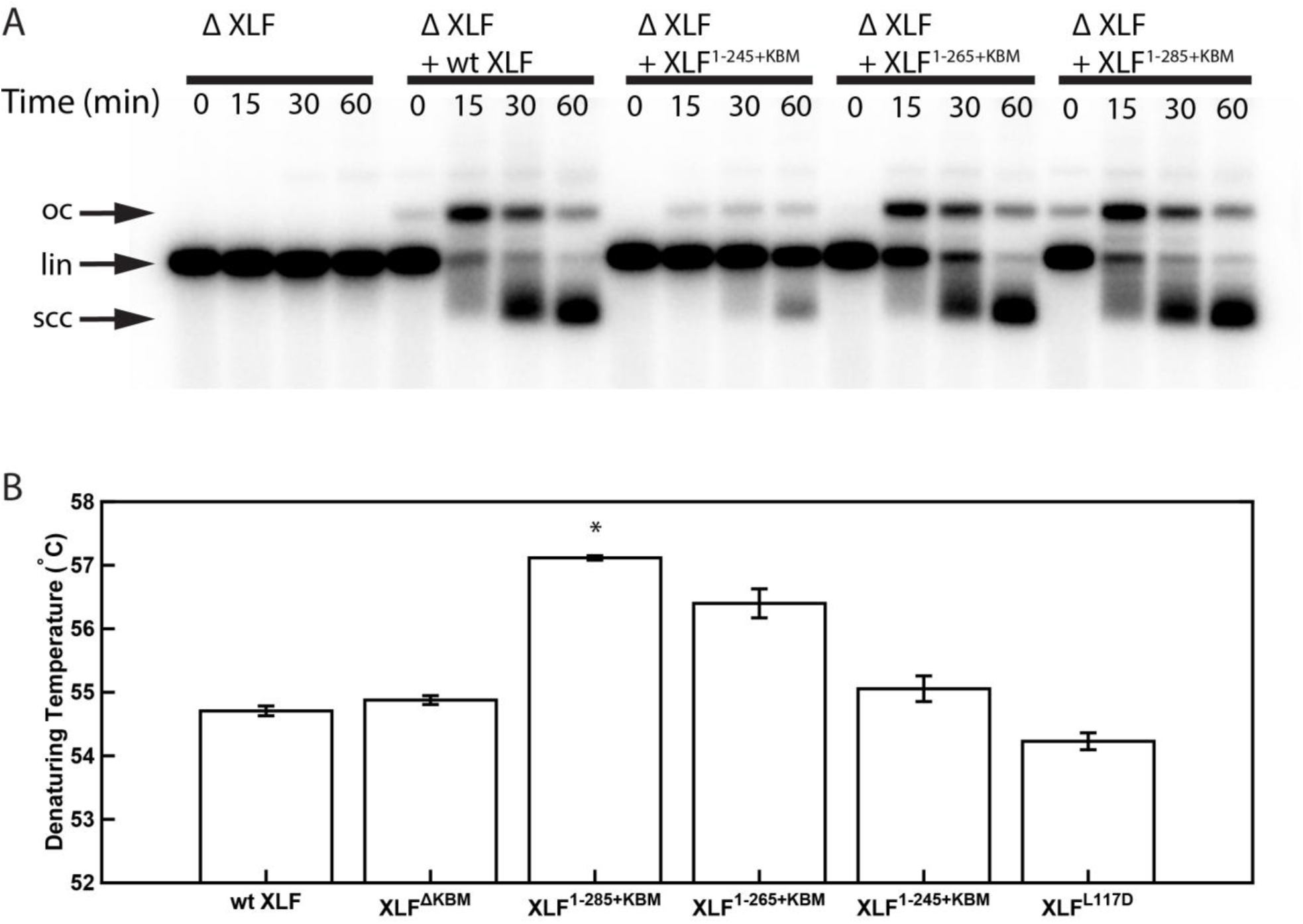
Characterization of XLF truncation mutants. **A**, Ensemble time course end joining assay in XLF-depleted *Xenopus* egg extract. Recombinant wt XLF, XLF^1-285+KBM^, XLF^1-265+KBM^, and XLF^1-245+KBM^ were added back at to their respective reaction samples at 500 nM final concentration. DNA species: scc, supercoiled closed circular; lin, linear; oc, open circle. **B**, Denaturing temperatures of XLF constructs as measured by differential scanning fluorimetry. The average of two replicates is shown with error bars representing the minimum and maximum. Only XLF^1-285+KBM^ had a denaturing temperature statistically different from wt XLF (p=0.0066), while the denaturing temperature of the other constructs (XLF^ΔKBM^ p = 0.2451, XLF^1-265+KBM^ p = 0.615, XLF^1-245+KBM^ p = 0.3091, XLF^L117D^ p = 0.1172) were not statistically different from that of wt XLF as determined by a two-tailed, unpaired *t* test with unequal variance and the Bonferroni correction. The range of denaturing temperatures was less than 3°C, suggesting no substantial changes in protein folding or stability.

**Supplementary Figure 3.**
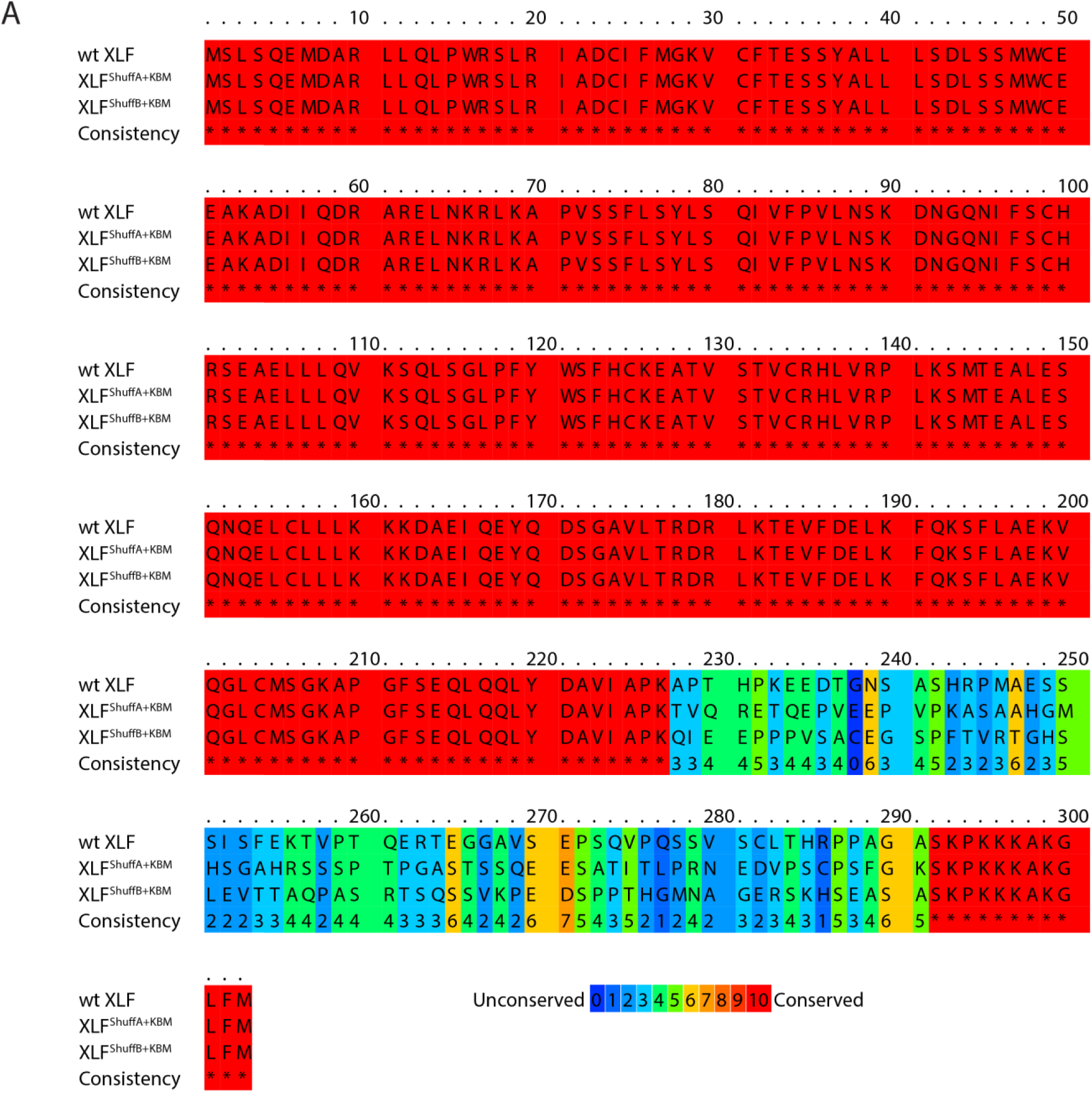
Sequence alignment of XLF shuffled tail mutants. **A**, Protein sequence alignment of wt XLF, XLF^ShuffA+KBM^, and XLF^ShuffB+KBM^. The alignment was performed using PRALINE multiple sequence alignment.^43^ Gap penalties were altered to prevent gaps in the alignment.

**Supplementary Figure 4.**
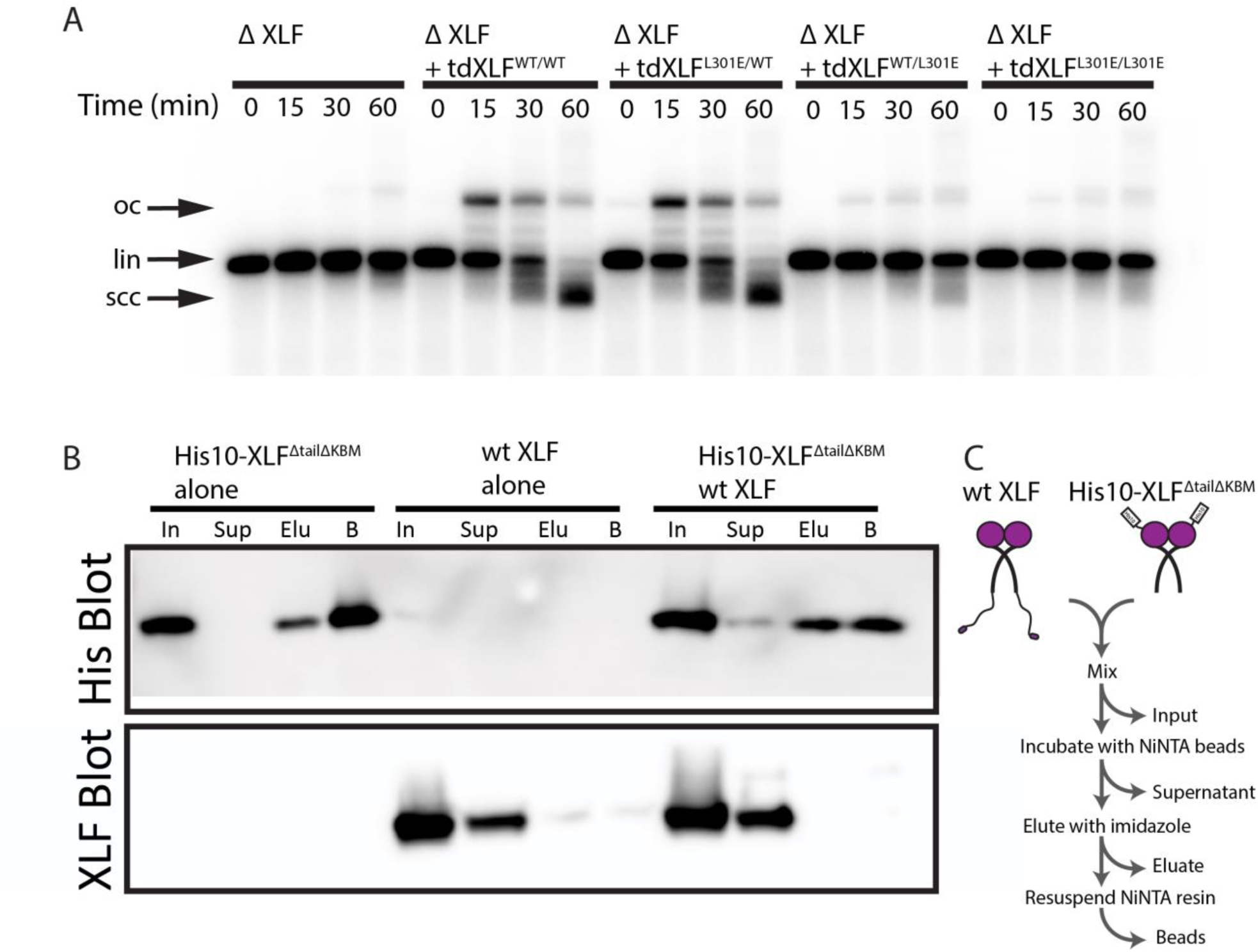
Characterization of the tdXLF and XLF heterodimer constructs. **A**, Ensemble time course end joining assay in XLF-depleted *Xenopus* egg extract. Recombinant tdXLF^WT/WT^, tdXLF^L301E/WT^, tdXLF^WT/L301E^, and tdXLF^L301E/L301E^ were added back at 50 nM final concentration to their respective reaction samples. DNA species: scc, supercoiled closed circular; lin, linear; oc, open circle. **B**, XLF heterodimer subunit exchange assay. His10-XLF^ΔtailΔKBM^ and wt XLF each alone or in combination were incubated at room temperature (input: I) before being incubated with NiNTA resin that was subsequently, spun down (supernatant: Sup), washed, and eluted (eluate: Elu), and resuspended (beads: B). Exchange between dimer subunits in the His10-XLF^ΔTailΔKBM^ + wt XLF sample would result in detecting wt XLF in the eluate and/or bead sample(s). **C**, Cartoon schematic of experimental workflow of the heterodimer exchange assay.

**Supplementary Figure 5.**
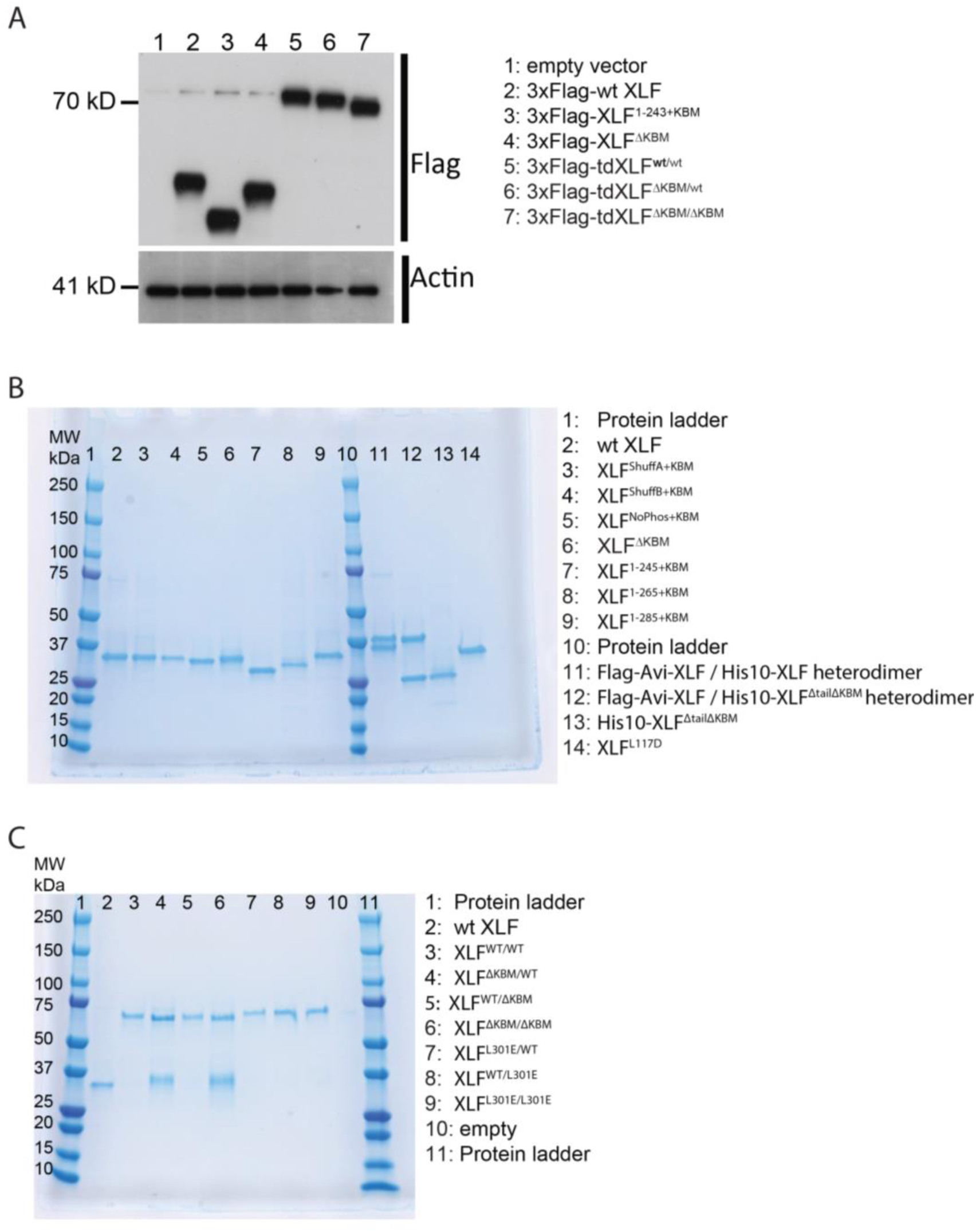
XLF constructs. **A**, Expression of XLF constructs in mESCs after transfection with the indicated constructs within the pCAGGS-BSKX vector. Samples of extract from each transfection were blotted with both anti-Flag and anti-Actin antibodies. **B**, XLF mutants and heterodimer constructs. 1500 ng of each construct was loaded on a 4-15% Mini-PROTEAN TGX Precast Protein Gel (BioRad). 3 µg of each heterodimer construct was loaded so that individual bands should be of the same intensity as the other XLF constructs. **C**, Tandem dimer XLF constructs. 750 ng of wtXLF and each tandem dimer construct were loaded and run on a 4-15% Mini-PROTEAN TGX Precast Protein Gel (BioRad). tdXLF constructs where the KBM of subunit 1 was replaced with additional linker sequence (lane 4 tdXLF^ΔKBM/WT^ and lane 6 tdXLF^ΔKBM/ΔKBM^) purified with a contaminant(s) that runs at ∼35 kDa on these PAGE gels. These contaminant bands were sent for mass spectrometry (Taplin Mass Spectrometry Core Facility, Harvard Medical School). Although peptides corresponding to *Xenopus* XLF were detected in both cases, the majority of peptides identified were from endogenous E.coli proteins, with the bacterial lipid synthesis protein, lpxD, being responsible for 72% and 41% of peptides identified from the contaminant bands within the tdXLF^ΔKBM/WT^ and tdXLF^ΔKBM/ΔKBM^ samples, respectively.

**Supp. Table 1.**

| **<u>TOTALS</u>** | **<u># Replicates</u>** | **<u># Molecules</u>** | **<u>Event #</u>** | **<u>Ave Formation Rate (s<sup>-1</sup>)</u>** | **<u>SEM (s<sup>-1</sup>)</u>** |
| --- | --- | --- | --- | --- | --- |
| Buffer | 3 replicates | 778 | 15 | 6.4x10<sup>-5</sup> | 1.0x10<sup>-5</sup> |
| wt XLF | 3 replicates | 691 | 267 | 1.7x10<sup>-3</sup> | 3.4x10<sup>-4</sup> |
| XLF<sup>1-245+KBM</sup> | 4 replicates | 1076 | 13 | 4.6x10<sup>-5</sup> | 1.3x10<sup>-5</sup> |

**Supp. Table 2.**
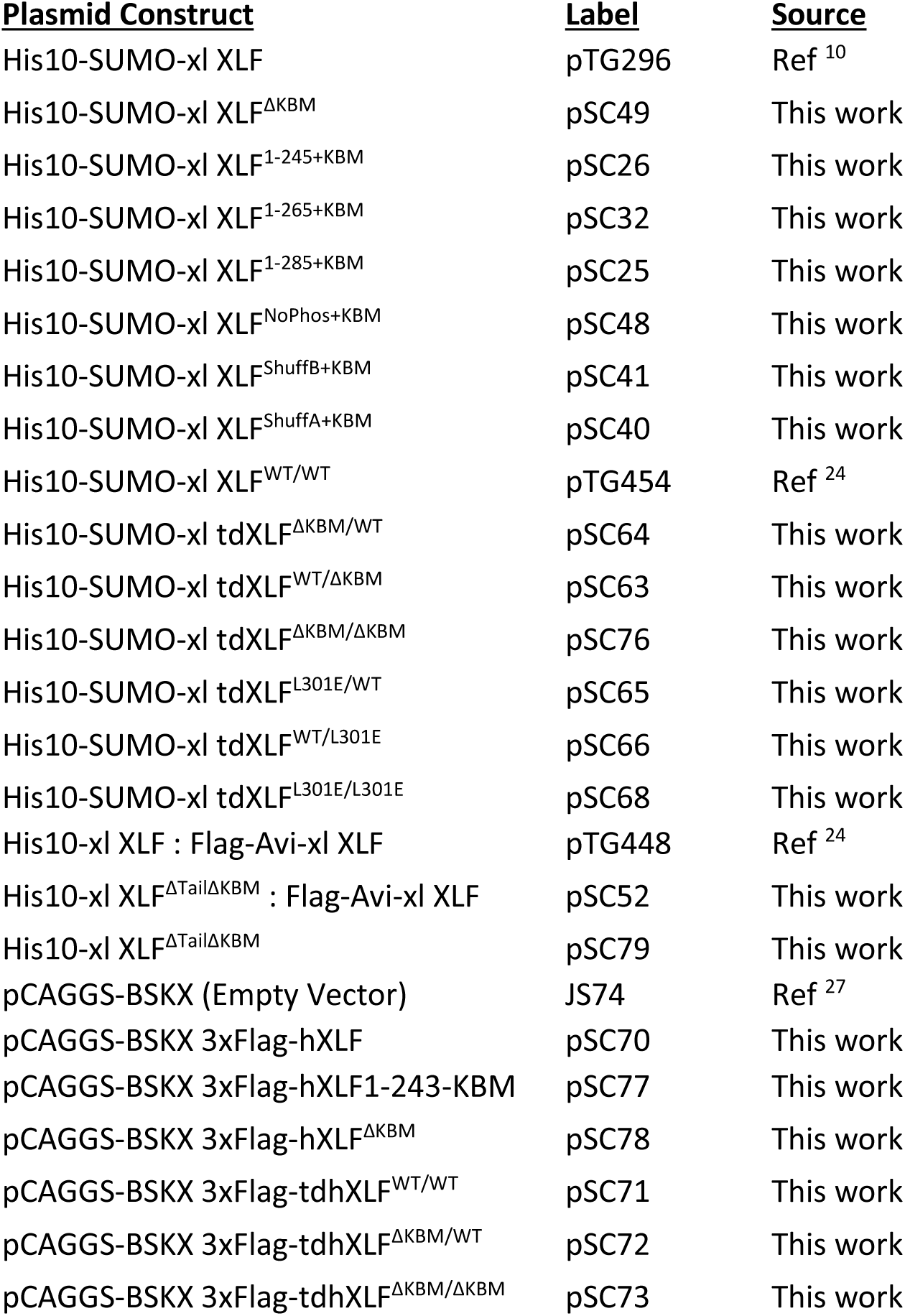

| **<u>Plasmid Construct</u>** | **<u>Label</u>** | **<u>Source</u>** |
| --- | --- | --- |
| His10-SUMO-xl XLF | pTG296 | Ref <sup>10</sup> |
| His10-SUMO-xl XLF<sup>ΔKBM</sup> | pSC49 | This work |
| His10-SUMO-xl XLF<sup>1-245+KBM</sup> | pSC26 | This work |
| His10-SUMO-xl XLF<sup>1-265+KBM</sup> | pSC32 | This work |
| His10-SUMO-xl XLF<sup>1-285+KBM</sup> | pSC25 | This work |
| His10-SUMO-xl XLF<sup>NoPhos+KBM</sup> | pSC48 | This work |
| His10-SUMO-xl XLF<sup>ShuffB+KBM</sup> | pSC41 | This work |
| His10-SUMO-xl XLF<sup>ShuffA+KBM</sup> | pSC40 | This work |
| His10-SUMO-xl XLF<sup>WT/WT</sup> | pTG454 | Ref <sup>24</sup> |
| His10-SUMO-xl tdXLF<sup>ΔKBM/WT</sup> | pSC64 | This work |
| His10-SUMO-xl tdXLF<sup>WT/ΔKBM</sup> | pSC63 | This work |
| His10-SUMO-xl tdXLF<sup>ΔKBM/ΔKBM</sup> | pSC76 | This work |
| His10-SUMO-xl tdXLF<sup>L301E/WT</sup> | pSC65 | This work |
| His10-SUMO-xl tdXLF<sup>WT/L301E</sup> | pSC66 | This work |
| His10-SUMO-xl tdXLF<sup>L301E/L301E</sup> | pSC68 | This work |
| His10-xl XLF : Flag-Avi-xl XLF | pTG448 | Ref <sup>24</sup> |
| His10-xl XLF<sup>ΔTailΔKBM</sup> : Flag-Avi-xl XLF | pSC52 | This work |
| His10-xl XLF<sup>ΔTailΔKBM</sup> | pSC79 | This work |
| pCAGGS-BSKX (Empty Vector) | JS74 | Ref <sup>27</sup> |
| pCAGGS-BSKX 3xFlag-hXLF | pSC70 | This work |
| pCAGGS-BSKX 3xFlag-hXLF1-243-KBM | pSC77 | This work |
| pCAGGS-BSKX 3xFlag-hXLF<sup>ΔKBM</sup> | pSC78 | This work |
| pCAGGS-BSKX 3xFlag-tdhXLF<sup>WT/WT</sup> | pSC71 | This work |
| pCAGGS-BSKX 3xFlag-tdhXLF<sup>ΔKBM/WT</sup> | pSC72 | This work |
| pCAGGS-BSKX 3xFlag-tdhXLF<sup>ΔKBM/ΔKBM</sup> | pSC73 | This work |

## References

1. Rothkamm, K., Krüger, I., Thompson, L.H., & Löbrich, M. Pathways of DNA Double-Strand Break Repair during the Mammalian Cell Cycle. Mol. Cell. Biol. 23, No. 16, 5706–5715 (2003).

2. Walker, J. R., Corpina, R. A. & Goldberg, J. Structure of the Ku heterodimer bound to dna and its implications for double-strand break repair. Nature 412, 607–614 (2001).

3. Uematsu, N. et al. Autophosphorylation of DNA-PKCS regulates its dynamics at DNA double-strand breaks. J. Cell Biol. 177, 219–229 (2007).

4. Jette, N. & Lees-Miller, S. P. The DNA-dependent protein kinase: A multifunctional protein kinase with roles in DNA double strand break repair and mitosis. Prog. Biophys. Mol. Biol. 117, 194–205 (2015).

5. Jiang, W. et al. Differential phosphorylation of DNA-PKcs regulates the interplay between end-processing and end-ligation during nonhomologous end-joining. Mol. Cell 58, 172–185 (2015).

6. Grawunder, U. et al. Activity of DNA ligase IV stimulated by complex formation with XRCC4 protein in mammalian cells. Nature 388, 492–495 (1997).

7. Stinson, B. M., Moreno, A. T., Walter, J. C. & Loparo, J. J. A Mechanism to Minimize Errors during Non-homologous End Joining. Mol. Cell 77, 1080-1091.e8 (2020).

8. Buck, D. et al. Cernunnos, a novel nonhomologous end-joining factor, is mutated in human immunodeficiency with microcephaly. Cell 124, 287–299 (2006).

9. Ahnesorg, P., Smith, P. & Jackson, S. P. XLF interacts with the XRCC4-DNA Ligase IV complex to promote DNA nonhomologous end-joining. Cell 124, 301–313 (2006).

10. Graham, T. G. W., Walter, J. C. & Loparo, J. J. Two-Stage Synapsis of DNA Ends during Non-homologous End Joining. Mol. Cell 61, 850–858 (2016).

11. Cottarel, J. et al. A noncatalytic function of the ligation complex during nonhomologous end joining. J. Cell Biol. 200, 173–186 (2013).

12. Wang, J. L. et al. Dissection of DNA double-strand-break repair using novel single-molecule forceps. Nat. Struct. Mol. Biol. 25, 482–487 (2018).

13. Xing, M. et al. Interactome analysis identifies a new paralogue of XRCC4 in non-homologous end joining DNA repair pathway. Nat. Commun. 6, (2015).

14. Ochi, T. et al. PAXX, a paralog of XRCC4 and XLF, interacts with Ku to promote DNA double-strand break repair. Science (80-.). 347, 185–188 (2015).

15. Li, Y. et al. Crystal structure of human XLF/Cernunnos reveals unexpected differences from XRCC4 with implications for NHEJ. EMBO J. 27, 290–300 (2008).

16. Andres, S. N., Modesti, M., Tsai, C. J., Chu, G. & Junop, M. S. Crystal Structure of Human XLF: A Twist in Nonhomologous DNA End-Joining. Mol. Cell 28, 1093–1101 (2007).

17. Andres, S. N. et al. A human XRCC4-XLF complex bridges DNA. Nucleic Acids Res. 40, 1868–1878 (2012).

18. Malivert, L. et al. Delineation of the Xrcc4-interacting region in the globular head domain of cernunnos/XLF. J. Biol. Chem. 285, 26475–26483 (2010).

19. Roy, S. et al. XRCC4/XLF Interaction Is Variably Required for DNA Repair and Is Not Required for Ligase IV Stimulation. Mol. Cell. Biol. 35, 3017–3028 (2015).

20. Roy, S. et al. XRCC4’s interaction with XLF is required for coding (but not signal) end joining. Nucleic Acids Res. 40, 1684–1694 (2012).

21. Ropars, V. et al. Structural characterization of filaments formed by human Xrcc4-Cernunnos/XLF complex involved in nonhomologous DNA end-joining. Proc. Natl. Acad. Sci. U. S. A. 108, 12663–12668 (2011).

22. Mahaney, B. L., Hammel, M., Meek, K., Tainer, J. A. & Lees-Miller, S. P. XRCC4 and XLF form long helical protein filaments suitable for DNA end protection and alignment to facilitate DNA double strand break repair. Biochem. Cell Biol. 91, 31–41 (2013).

23. Reid, D. A. et al. Organization and dynamics of the nonhomologous end-joining machinery during DNA double-strand break repair. Proc. Natl. Acad. Sci. U. S. A. 112, E2575–E2584 (2015).

24. Graham, T. G. W., Carney, S. M., Walter, J. C. & Loparo, J. J. A single XLF dimer bridges DNA ends during nonhomologous end joining. Nat. Struct. Mol. Biol. 25, 877–884 (2018).

25. Yano, K. I., Morotomi-Yano, K., Lee, K. J. & Chen, D. J. Functional significance of the interaction with Ku in DNA double-strand break recognition of XLF. FEBS Lett. 585, 841–846 (2011).

26. Nemoz, C. et al. XLF and APLF bind Ku80 at two remote sites to ensure DNA repair by non-homologous end joining. Nat. Struct. Mol. Biol. 25, 971–980 (2018).

27. Bhargava, R. et al. C-NHEJ without indels is robust and requires synergistic function of distinct XLF domains. Nat. Commun. 9, (2018).

28. Yu, Y. et al. DNA-PK and ATM phosphorylation sites in XLF/Cernunnos are not required for repair of DNA double strand breaks. DNA Repair (Amst). 7, 1680–1692 (2008).

29. Wühr, M. et al. Deep proteomics of the xenopus laevis egg using an mRNA-derived reference database. Curr. Biol. 24, 1467–1475 (2014).

30. Graham, T. G. W., Walter, J. C. & Loparo, J. J. Ensemble and Single-Molecule Analysis of Non-Homologous End Joining in Frog Egg Extracts. Methods Enzymol. 591, 233–270 (2017).

31. Stothard, P. The Sequence Manipulation Suite. Biotechniques 28, 1102–1104 (2000).

32. Wu, P. Y. et al. Interplay between Cernunnos-XLF and nonhomologous end-joining proteins at DNA ends in the cell. J. Biol. Chem. 282, 31937–31943 (2007).

33. Frit, P., Ropars, V., Modesti, M., Charbonnier, J. B. & Calsou, P. Plugged into the Ku-DNA hub: The NHEJ network. Prog. Biophys. Mol. Biol. (2019). doi:10.1016/j.pbiomolbio.2019.03.001

34. Hammel, M. et al. XRCC4 protein interactions with XRCC4-like factor (XLF) create an extended grooved scaffold for DNA ligation and double strand break repair. J. Biol. Chem. 286, 32638–32650 (2011).

35. Malivert, L. et al. The C-Terminal Domain of Cernunnos/XLF Is Dispensable for DNA Repair In Vivo. Mol. Cell. Biol. 29, 1116–1122 (2009).

36. Normanno, D. et al. Mutational phospho-mimicry reveals a regulatory role for the XRCC4 and XLF C-terminal tails in modulating DNA bridging during classical non-homologous end joining. Elife 6, 1–28 (2017).

37. Zhao, B. et al. The essential elements for the noncovalent association of two DNA ends during NHEJ synapsis. Nat. Commun. 10, 3588 (2019).

38. Hemsley, A., Arnheim, N., Toney, M. D., Cortopassi, G. & Galas, D. J. A simple method for site-directed mutagenesis using the polymerase chain reaction. Nucleic Acids Res. 17, 6545–6551 (1989).

39. Gibson, D. G. et al. Enzymatic assembly of DNA molecules up to several hundred kilobases. Nat. Methods 6, 343–345 (2009).

40. Lebofsky R., Takahashi T., and W. J. C. DNA Replication in Nucleus-Free Xenopus Egg Extracts. In Methods in Molecular Biology 521, 229–252 (2009).

41. Lebofsky, R., Van Oijen, A. M. & Walter, J. C. DNA is a co-factor for its own replication in Xenopus egg extracts. Nucleic Acids Res. 39, 545–555 (2011).

42. Campbell, R. notBoxPlot. GitHub. Retrieved from https://github.com/raacampbell/notBoxPlot on April 24 2018 (2020).

43. Simossis, V. A. & Heringa, J. PRALINE: A multiple sequence alignment toolbox that integrates homology-extended and secondary structure information. Nucleic Acids Res. 33, 289–294 (2005).

